# Identification of pan-cancer/testis genes and validation of therapeutic targeting in triple-negative breast cancer: Lin28a- and Siglece-based vaccination induces anti-tumor immunity and inhibits metastasis

**DOI:** 10.1101/2023.05.09.539617

**Authors:** Jason A. Carter, Bharati Matta, Jenna Battaglia, Carter Somerville, Benjamin D. Harris, Margaret LaPan, Gurinder S. Atwal, Betsy J. Barnes

## Abstract

**Background:** Cancer-testis (CT) genes are targets for tumor antigen-specific immunotherapy given that their expression is normally restricted to the immune-privileged testis in healthy individuals with aberrant expression in tumor tissues. While they represent targetable germ-tissue antigens and play important functional roles in tumorigenesis, there is currently no standardized approach for identifying clinically relevant CT genes. Optimized algorithms and validated methods for accurate prediction of reliable CT antigens with high immunogenicity are also lacking.

**Methods:** Sequencing data from the Genotype-Tissue Expression (GTEx) and The Genomic Data Commons (GDC) databases was utilized for the development of a bioinformatic pipeline to identify CT exclusive genes. A CT germness score was calculated based on the number of CT genes expressed within a tumor type and their degree of expression. The impact of tumor germness with clinical outcome was evaluated using healthy GTEx and GDC tumor samples. We then used a triple-negative breast cancer mouse model to develop and test an algorithm that predicts epitope immunogenicity based on the identification of germline sequences with strong MHCI and MHCII binding affinities. Germline sequences for CT genes were synthesized as long synthetic peptide vaccines and tested in the 4T1 triple-negative model of invasive breast cancer with Poly(I:C) adjuvant. Vaccine immunogenicity was determined by flow cytometric analysis of *in vitro* and *in vivo* T cell responses. Primary tumor growth and lung metastasis was evaluated by histopathology, flow cytometry and colony formation assay.

**Results:** We developed a new bioinformatic pipeline to reliably identify CT exclusive genes as immunogenic targets for immunotherapy. We identified CT genes that are exclusively expressed within the testis, lack detectable thymic expression, and are significantly expressed in multiple tumor types. High tumor germness correlated with tumor progression but not with tumor mutation burden, supporting CT antigens as appealing targets in low mutation burden tumors. Importantly, tumor germness also correlated with markers of anti-tumor immunity. Vaccination of 4T1 tumor bearing mice with Siglece and Lin28a antigens resulted in increased T cell anti-tumor immunity and reduced primary tumor growth and lung metastases.

**Conclusion:** Our results present a novel strategy for the identification of highly immunogenic CT antigens for the development of targeted vaccines that induce anti-tumor immunity and inhibit metastasis.

## Introduction

Immunotherapy relies on the ability of the adaptive immune system to differentiate neoplastic cells from healthy tissues, most commonly via the recognition of tumor neoantigens arising from somatic mutations^1, 2^. Personalized cancer vaccines enable targeting of specific tumor neoantigens^3, 4^ and may therefore work synergistically with immune checkpoint blockade (ICB) therapies^5^, with vaccination against immunogenic mutations in melanoma^6–8^ and glioblastoma^9–11^ patients showing early clinical promise. However, random somatic mutations, aptly referred to as private neoantigens, are rarely shared between individuals and therefore have limited generalizability across patients^12^. Public neoantigens shared across many tumors, in contrast, have off-the-shelf therapeutic potential but have been largely restricted to rare cancer driver mutations^13^. Cancer-testis (CT) genes represent a distinct alternative source of cancer related antigens potentially amenable to a broad class of immunotherapies, include personalized vaccines and cell therapy.

CT genes, defined by their expression restricted to immune-privileged testis and ectopic expression in cancer, contain germline sequences that are immunogenic when ectopically expressed by neoplastic cells^14–16^. While the aberrant expression of CT genes has historically thought to arise from epigenetic dysregulation^17, 18^, recent work has increasingly demonstrated crucial roles for CT gene expression in oncogenesis^19^, metastatic disease progression^20–22^ and potentially ICB response^23^. CT gene upregulation has been observed across numerous tumor types^24^, including those with relatively low tumor mutational burdens such as hormone-receptor negative tumors^25–27^, making them a potentially widely applicable target for new immunotherapeutics. Despite the promising results of CT antigen-based immunotherapy in many cancers such as melanoma, glioblastoma, and ovarian tumors (NCT02166905; NCT00071981; NCT01213472; NCT02650986; NCT01967823), such vaccination trials in breast cancer patients are still limited or get withdrawn due to null findings (NCT03110445; NCT01522820; NCT03093350; NCT02015416)^26^.

The CTdatabase provides a repository of over 200 known CT genes compiled from the literature ^28^. However, the methods used in CT gene discovery and even the criteria used to define CT gene expression thresholds have not been standardized^29–31^. The development of the Genotype-Tissue Expression (GTEx)^32^ and the Genomic Data Commons (GDC), including The Cancer Genome Atlas (TCGA)^33^ and Therapeutically Applicable Research to Generate Effective Treatments (TARGET)^33, 34^, public sequencing repositories for various healthy tissues and cancer types, respectively, have more recently enabled high-throughput screening for CT expression patterns^35, 36^. These approaches have not focused on the identification of immunogenic CT genes for targeted vaccine therapies, however, which requires a high degree of testis exclusivity, a lack of central tolerance (*i.e.*, absent thymic expression), and robust cancer expression that ideally represents function as an oncogenic driver.

We here implemented a new bioinformatic pipeline to perform an unbiased, genome-wide screen aimed at identifying genes with robust CT expression that are more likely to represent immunogenic targets for immunotherapy. We identified 103 CT genes, 70 (68%) of which are not present in the CTdatabase, that meet a highly conservate exclusivity expression threshold, lack detectable thymic expression, and are significantly expressed in multiple tumor types. We further developed and validated the use of an *ad hoc* algorithm that incorporates both MHCI and MHCII antigen binding strength predictions^37^ to optimize selection of immunogenic CT gene germline sequences for synthetic long peptide vaccination^38^. Two test CT genes – *Siglece* and *Lin28a* – were identified for validation based on their testis-exclusive expression in humans and their differential expression in 4T1 mouse primary mammary tumors and lung metastases. CT antigen vaccination against Siglece or Lin28a resulted in a robust CD4+ T cell helper and CD8+ T cell cytolytic response, with a significant reduction in primary tumor growth and lung metastases in a mouse model of triple-negative breast cancer.

## Methods

### Identification of testis-exclusive genes

Gene level RNA-seq data was downloaded directly from version 8 of the Genotype-Tissue Expression (GTEx) project^32^. In all, we obtained RNA-sequencing data from 17,382 healthy samples representing 30 different tissue types with gene expression quantified as transcripts per million (TPM). We then used the Ensembl^39^ annotation to identify those genes which were protein coding, removing those without protein IDs from downstream analysis. To differentiate tissue samples which actively expressed each gene from transcriptional and technical noise, we chose a threshold of 2 TPM^40^. Using this conservative threshold, we then calculated the Phi Correlation Coefficient^41^ (φ) for each protein-coding gene with respect to all 30 GTEx healthy tissue types. Specfically, φ_*i*_for the i^th^ tissue type and a given gene is calculated as:

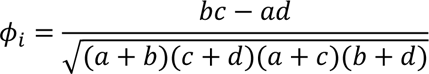

where a is the number of i^th^ tissue samples expressing that gene (at a threshold of ≥ 2 TPM), b is the number of samples with gene expression across all other tissue types, c is the number of i^th^ tissue samples lacking expression of that gene, and d is the number of all other tissue samples without gene expression. A φ value of 0.95 corresponds to approximately 2.5% of all samples having non-exclusive expression (i.e. 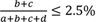) Gene ontology analysis was conducted using th.e PANTHER classification system^42^.

### Identification of genes without thymic expression

RNA-sequencing data from human medullary thymic epithelial cell (mTEC) samples was obtained under accession codes GSE201720^43^ and GSE127209^44^. We included both major histocomabitility complex class II (MHCII) low (immature) and high (mature) mTEC samples from both previously published datasets. Transcript abundance quantification was performed using Kallisto^45^ and a threshold of 2 TPM was again used a threshold for thymic expression. Testis-exclusive genes with expression in any thymic sample were excluded from further analysis.

### Identification of cancer/testis genes

We next examined the expression of these protein-coding, testis-exclusive genes without thymic expression across 13,237 tumor samples representing 38 cancer types. Tumor sequencing data and clinical data was obtained from the GDC^46^, including both TCGA and the TARGET initiative^33, 34^. Cancer types with at sequencing data for at least 50 samples were included. Using the expression threshold of 2 TPM, we calculated the fraction of tumors in each cancer type expressing each gene. We defined cancer/testis gene expression as those testis-exclusive genes without thymic expression that were expressed in at least 5% of tumors across two different tumor types. Given their high expression of testis-exclusive genes, testicular germ cell tumors (TGCT) were not included in the determination of cancer/testis gene expression. Previously identified CT genes were obtained from CTdatabase^28^.

### Evaluation of germness score

We defined a germness score as the average expression of our 103 CT genes, explicitly:

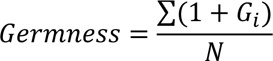

where *G*_*i*_ is the expression (in TPM) of the *i*^th^ CT gene and N is the total number of CT genes. We calculated the germness score for each GTEx healthy and GDC tumor sample. Clinical data was obtained from GDC^46^ and additional features of the tumor immune microenvironment for GDC samples were obtained from Thorsson *et al.*^47^. Kaplan-Meier curves were generated using the Python lifelines package^48^.

### Mice

Wild-type Balb/c mice were purchased from The Jackson Laboratory (Bar Harbor, ME). Due to the nature of the study, only 8-12 weeks-old female mice were used. Mice were kept in environment-controlled pathogen–free conditions with a 12-hour light/dark cycle and an ambient temperature of 23°C ± 2°C. This study was carried out in strict accordance with recommendations in *Guide for the Care and Use of Laboratory Animals* of the NIH (National Research Council, Guide for the Care and Use of Laboratory Animals (National Academies Press, Washington, DC, ed. 8, 2011). The protocol was approved by the Institutional Animal Care and Use Committee (IACUC) of the Feinstein Institutes for Medical Research.

### Selection of CT antigen sequences for vaccination

We sequenced 6 replicates from healthy thymus, mammary, and lung mouse tissue samples, as well as thymic samples from tumor-bearing mice, primary mammary tumors, and lung metastases. Abundance quantification and differential expression analyses were performed using Kallisto^45^ and Sleuth^49^, respectively. After examining expression of mouse homologs of human protein-coding, testis-exclusive genes without thymic expression across each of these 36 healthy and tumor-bearing mouse samples, we identified *Lin28a* and *Siglece* as potential CT vaccine targets due to expression limited to primary mammary tumors and/or lung metastases.

The entire germline sequences of both *Lin28a* and *Siglece* were divided into all possible 25-30 amino acid substrings and the number of peptides predicted to bind to H-2-Dd, H-2-Kd, H-2-Ld, H-2-IAd and H-2-IEd calculated for each substring using NetMHCpan^37^. Each sequence was then scored according to the number of strong (top 0.5% and 1% of predicted MHCI and MHCII binding) and weak (top 2% and 5%, respectively) binders, with weak binders weighted as ¼ of strong binders. The two top scoring distinct peptide sequences were selected for both Siglece (S) and Lin28a (L). Amino acid sequences chosen for synthesis were: KKDAGLYFFRLERGKTKYNYMWDKMTLVV (S1), TRMTIRLNVSYAPKNLTVTIYQGADSVSTI (S2), KLPPQPKKCHFCQSINHMVASCPLKAQQGP (L1), and EAVEFTFKKSAKGLESIRVTGPGGVFCIGS (L2).

### In vivo tumor initiation and metastasis

To initiate primary tumor growth and lung metastasis, freshly prepared 1 x 10^5^ 4T1 cells in mid-log phase growth were orthotopically injected into the 4^th^ right mammary fat pad of Balb/c mice. Two vaccination protocols were assessed – 1) pre- and 2) post-tumor cell inoculation. Tumor cell inoculation day was determined as day 0 on the dosing strategy chart. Mice were sacrificed on different days depending on the experiment, tumors weighed, and organs harvested for fixation and histopathological analysis, flow cytometry analysis of T cell composition and T cell proliferation assay. Lung metastases were quantified by counting the number of visible nodules on the surface of the lung. Counting was performed in a blinded manner by 3 independent readers.

### Lung clonogenic assay

Mice from the prevention strategy were euthanized on day 28 and lungs were harvested from treated mice. Single cell suspension was made, and 200 cells were plated in complete media with 60 µM 6-Thioguanine for 14 days. Non-adherent cells were washed out, adherent 4T1 clones were fixed with methanol for an hour, washed and stained with 0.5% crystal violet for 2 hours followed by washing with PBS. Images were taken on Evos M700 imaging system. Each dot represents a 4T1 colony, the dots were counted for quantitation.

### Splenocyte proliferation assay

Spleens of vaccinated mice (S1, L1 or L2; 100µg) were harvested 2 weeks after the last injection and single cell suspension was made. The cells were stained with 5μm CFSE (Invitrogen:C34554). Briefly, cells were incubated in dark with the dye for 8-10 minutes. The cells were then washed twice with 10ml ice cold PBS+10%FBS to get rid of excess dye. The cells were then plated along with matched antigen (S1, L1 or L2; 1μg/well) for 4 days. Similarly for controls, CFSE stained splenocytes from naïve mice were also plated with all three peptides in culture. On day 5, cells were collected and stained with surface markers for T cells. The samples were run using BD Fortessa and analyzed on BD FlowJo and ModFit.

### Flow cytometry

Spleens and tumors were collected, and single cell suspension was made. The cells were then treated with RBC lysis buffer, washed, and blocked in PBS supplemented with Fc Blocker (BioLegend #422302) for 15 min and then stained with antibodies against surface makers for 30 minutes. After staining, cells were washed two times in PBS without Mg^++^ or Ca^2+^ and then fixed in 4% PFA before analysis on a BD Fortessa. For intracellular staining, after fixation, cells were permeabilized in 0.1% Triton X-100 followed by staining with intracellular markers. Most of the flow cytometry antibodies were purchased from BioLegend (CD45 PerCP#103130, CD3-BV650#100229, CD4-BV421#100443, CD8-PerCP#100732, CD8-PE#140408, IFNγ-APC#505809, Granzyme B-FITC#515403). Live/dead fixable stain Aqua #L34957, Green #L23105 were obtained from Thermo Fisher Scientific.

### Statistical analysis and code availability

Custom analysis code was written in Python (version >3.8) or R (version >4.03) for the identification of CT genes using GTEx and GDC data. All boxplots show median values with interquartile ranges and extrema (whiskers at 1.5x interquartile range [IQR]). Comparisons between paired samples were performed using a paired t-test, otherwise statistical significance was assessed using a Mann-Whitney U test. Correction for multiple hypothesis testing was performed using Bonferonni correction. GraphPad Prism software was used for statistical analyses of the mouse vaccination studies. Quantitative data are presented as mean ± SEM. Non-parametric data were analyzed by two-tailed Mann–Whitney *U* tests. All the data were compared between the different immunization groups by One-way ANOVA through Tukey’s *post hoc* tests. *p* < 0.05 was considered statistically significant for all analyses.

## Results

### Identification of immunogenic cancer-testis genes

While the CTdatabase^28^ provides a resource of known genes with CT expression, many of these genes were identified in early studies utilizing only limited sample sizes. While more recent work has utilized large public sequencing repositories, there is no standardized criteria defining thresholds for CT expression, nor have these studies been focused on the high throughput identification of immunogenic CT genes. With this in mind, we first implemented a bioinformatic pipeline to identify robust CT gene expression from existing sequencing repositories spanning more than 30,000 healthy tissue and tumor samples (**Figure 1A**). As our goal was to identify genes with the highest probability of serving as immunogenic targets for standardized “off-the-shelf” immunotherapy, we (1) identified protein-coding genes with robust testis-exclusive expression, (2) removed those testis-exclusive genes that demonstrated any human thymic expression (and thereby are most likely to have central tolerance despite testis-exclusive peripheral expression), and (3) identified those, from this subset of testis-exclusive genes without detectable thymic expression, with meaningful expression in multiple tumor subtypes. Together, our high-throughput screening approach identified 103 high-confidence CT gene candidates, including 70 CT genes that are not present in the CTdatabase and are potentially public antigens.

**Figure 1.**
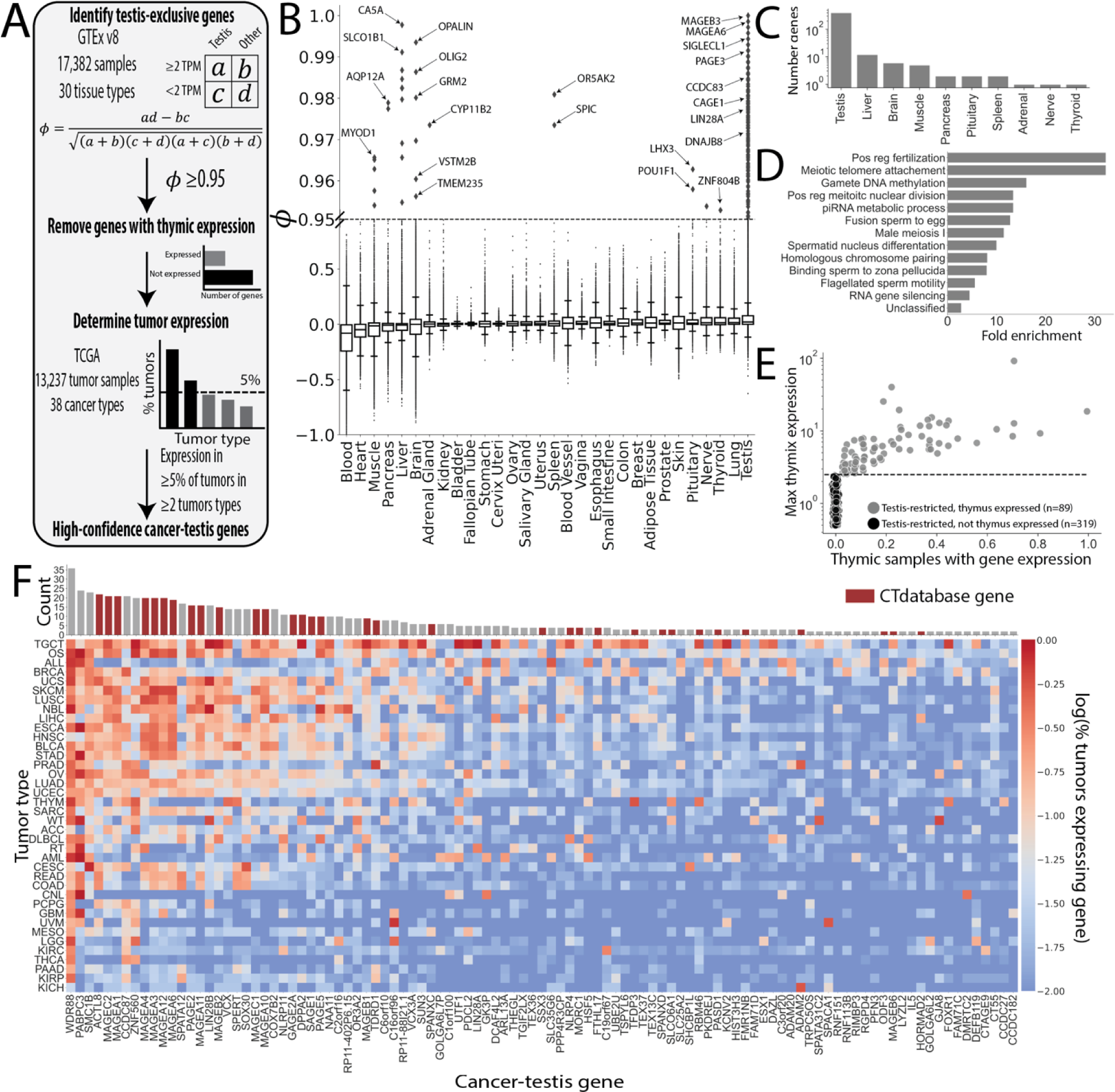
Identification of high-confidence cancer-testis genes. **(A)** Schematic overview of our bioinformatic pipeline to identify cancer-testis genes. Briefly, we identified testis-exclusive genes among GTEx healthy tissues using the Phi Correlation Coefficient^41^ (ϕ), excluded those with detectable thymic expression, and then identified a subset with robust tumor expression. **(B)** Distribution of gene exclusivity scores (ϕ) by GTEx tissue type. Those genes with ϕ≥0.95 are considered exclusive to that tissue type. **(C)** Number of tissue-exclusive genes found for each tissue type. **(D)** Gene ontology enrichment analysis^50, 51^ of testis-exclusive genes. Pos Reg-Positive regulation. **(E)** Thymic expression of testis-exclusive genes. Those testis-exclusive genes with detectable thymic expression (≥2 transcripts per million [TPM] in at least 1 thymic sample) were excluded. **(F)** The fraction of tumors expressing (≥2 TPM) a given testis-exclusive gene was calculated for each TCGA tumor type. CT genes were defined as those testis-exclusive genes without thymic expression that were expressed in at least 5% of tumors from at least 2 different tumor types.

Specifically, as CT genes are strictly defined by an expression pattern limited to healthy testis tissue with undetectable expression in all other tissue types, we first examined gene expression in 17,000 healthy samples from 30 tissue types in the GTEx database^32^. Using a threshold of 2 TPM to differentiate active gene expression from transcriptional noise^40^, we calculated the Phi Correlation Coefficient^41^ (φ) for each protein-coding gene in each tissue type (**Figure 1A**). To identify gene expression restricted to a single tissue type with a high-degree of confidence, we considered only genes with a φ greater than or equal to 0.95 in a particular tissue type to be tissue-exclusive (**Figure 1B**). Using this framework, we identified 440 tissue-exclusive protein-coding genes of which 408 (93%) were exclusively expressed in the testis (**Figure 1C** and **Supplemental Table 1&2**). Gene ontology analysis^50, 51^ of these testis-exclusive genes confirmed a range of testis-related functions including positive regulation of fertilization, gamete DNA methylation, and male meiosis I (**Figure 1D**).

As the testis are immune-privileged, testis-exclusive genes do not have the same central tolerance requirement as genes expressed in other peripheral tissues. Thus, since an antigen is targetable if and only if there is no central tolerance against that gene, we next directly examined thymic expression of these testis-exclusive genes to further maximize the chance of selecting genes with immunogenic germline sequences. We identified 89 of our 408 (22%) testis-exclusive genes that did have detectable thymic expression across in healthy human thymic samples^43, 44^, leaving 319 protein-coding testis-exclusive genes with undetectable thymic expression and thereby a lower chance of being recognized as a normal self-peptide (**Figure 1E** and **Supplemental Figure 1**). Of note, we found that only 64 out of 221 (29%) CTdatabase genes met our threshold for testis-exclusive expression, with an additional 8 of these 64 (12.5%) genes having detectable thymic expression (**Supplemental Figure 2**). Together, only 56 out of 221 (25%) CTdatabase genes met our conservative thresholds for CT gene expression and these known CT genes compose only 18% of our final 319 testis-exclusive genes without thymic expression (**Supplemental Table 2**).

We further examined expression of this set of testis-exclusive protein-coding genes lacking thymic expression across more than 13,000 individual tumor samples spanning 38 cancer types from the GDC, including data from both TCGA and the TARGET initiative. As we were interested in the identification of potential CTs that could represent widely conserved antigens, we screened our above set of testis-exclusive genes without thymic expression to identify CT genes that were expressed in at least 5% of tumors (again using an expression threshold of 2 TPM) in at least two different cancer types (**Figure 1F** and **Supplemental Figure 3**). Using this conservative definition, we identified 103 CT genes of which only 33 (32%) are present in the CTdatabase (**Supplemental Table 2**).

### CT gene expression correlates with tumor stage and anti-tumor immunity

We next explored CT gene expression across GDC tumor types. To facilitate comparison, we defined a germness score as the log average expression of our set of 103 CT genes in each sample, with higher germness scores correlating with more robust CT gene expression. We first compared germness between GDC tumor samples and paired healthy samples taken from adjacent parenchyma in the same individual. We found higher germness in tumor samples in 15 (10 stastistically significant) of 18 tumor types relative to paired normal samples (**Figure 2A**). Only kidney chromophobe (KICH) had significantly higher germness in adjacent healthy tissue. Further examination across tumor types and their corresponding GTEx healthy tissues demonstrated widespread upregulation of CT gene expression in this pan-cancer cohort (**Figure 2B**).

**Figure 2.**
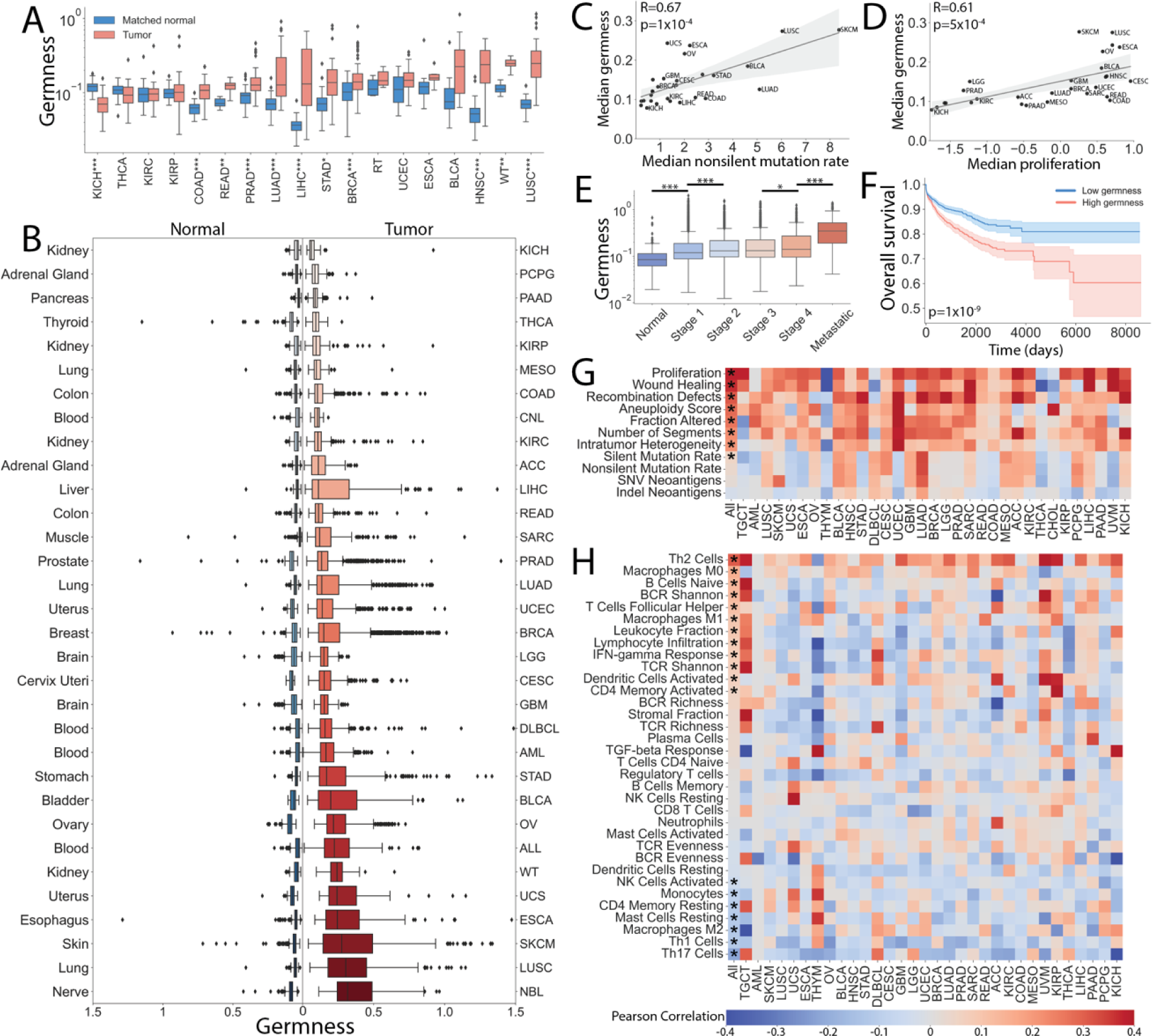
Association of cancer-testis gene expression and tumor characteristics. **(A)** Germness was defined as the log average expression of the 103 CT genes identified in this study. We found that comparison of germness between paired GDC tumor samples and tumor-adjacent healthy parenchymal samples in a majority of cancer types. *p ≤0.05, **p ≤0.01, ***p ≤0.001 by paired t-test test after Bonferroni correction. **(B)** Germness was then calculated for each GDC tumor type (with at least 50 samples) and it’s corresponding GTEx healthy tissue type, again demonstrating significantly higher germness in tumor samples relative to their respective healthy parenchyma. **(C)** Cancer types with higher non-silent mutation rates (R=0.67, *p*=1.1×10^−4^ by Wald test) and **(D)** proliferation rates (R=0.61, *p*=5.4×10^−4^) tended to have higher germness scores. **(E)** Increased germness scores are correlated with more advanced tumor stage. *p ≤0.05, ***p ≤0.001 by Mann-Whitney U test. **(F)** In the TCGA pan-cancer cohort, tumors with high germness (top 50%) had decreased overall survival rates relative to low germness tumors (p=1.1×10^−9^ by log rank test). **(G)** The Pearson correlation was calculated for tumor and **(H)** tumor-immune microenvironment features taken from Thorsson *et al.*^47^ Statistical significance (p ≤0.05 by Wald test after Bonferonni correction) across all tumor types is denoted by an asterisk (*).

Further examining CT gene expression across various cancer types, we found positive correlations between median tumor germness and non-silent mutation rate (**Figure 2C**, R=0.66, *p*=1.5 x 10^−4^ by Wald test), as well as between germness and tumor proliferation^52^ (**Figure 2D**, R=0.64, *p*=2.2 x 10^−4^ by Wald test). Consistent with CT gene expression being higher in cancer types with higher mutation rates and proliferation, we found that germness tended to increase with tumor stage in this pan-cancer cohort (**Figure 2E**), with germness significantly increasing between stage 1 and 2 (*p*=4.2 x 10^−7^ by Mann-Whitney U test) and between stage 3 and 4 of primary tumors (*p*=0.05 by Mann-Whitney U test). Further, consistent with a correlation between CT gene expression and the epithelial-mesenchymal transition^53^, we found that germness was significantly higher in metastatic tumor samples when compared to stage 4 primary tumors (**Figure 2E**, *p*=4.0 x 10^−50^ by Mann-Whitney U test). Similarly, as expected in more advanced disease states, overall survival was significantly higher in tumors with low germness scores (**Figure 2F**, *p*=1.1×10^−9^ by log rank test).

We next calculated the Pearson correlation between germness and various tumor features within each tumor type. We found strong correlations across tumor types for tumor proliferation, recombination defects, tumor aneuploidy, and intratumor heterogeneity (**Figure 2G**). While cancer types with higher rates of non-silent mutation tended to have higher germness scores, we did not identify a significant correlation between germness and non-silent mutation rate, single-nucleotide variant (SNV) neoantigens, or Indel neoantigens at the individual tumor level. Similarly comparing features of the tumor immune microenvironment, we found that germness was positively correlated with features including leukocyte fraction, IFN-γ response, Th2 cells, undifferentiated M0 and pro-inflammatory M1 macrophages, and T cell receptor (TCR) Shannon entropy (**Figure 2H**). Germness was additionally negatively correlated with the intra-tumoral presence of anti-inflammatory M2 macrophages, as well as memory CD4^+^, Th1, and Th17 T cells.

### Bioinformatic development of an immunogenic cancer-testis antigen peptide vaccine

We have so far identified a set of high-confidence testis-exclusive genes that are likely to contain immunogenic germline antigens when ectopically expressed in tumors (**Supplemental Table 2**). Further, we have shown that expression of these genes in human cancers is correlated with more advanced tumor stages and often with markers of anti-tumor immunity. To begin to explore the potential of these CT genes as specific targets for immunotherapy, we focused on the identification of CT antigens (CTAs) in breast cancer (**Supplemental Table 3**). We chose to focus on breast cancer since it represents a mid-to-low mutational burden cancer type, in contrast to the high mutational burden tumor types for which current immunotherapies have shown the greatest level of success. Moreover, we directed our attention toward triple-negative breast cancers (TNBCs, or basal-like tumors that lack ER, PR, and HER2 expression) as they represent 10–20% of all breast cancers, are highly aggressive, exhibit metastases, lack targeted therapies, and have a poor prognosis. TNBCs are not responsive to approved hormone therapies and thus therapeutic options remain limited. Consistent with previous studies showing higher CT gene expression in hormone-receptor negative tumors^54^, we found the highest germness scores for the basal-like breast cancer subtype amongst GDC samples^55, 56^ (**Figure 3A**). We therefore elected to use the 4T1 murine mammary cancer cell line as a model of TNBC.

**Figure 3.**
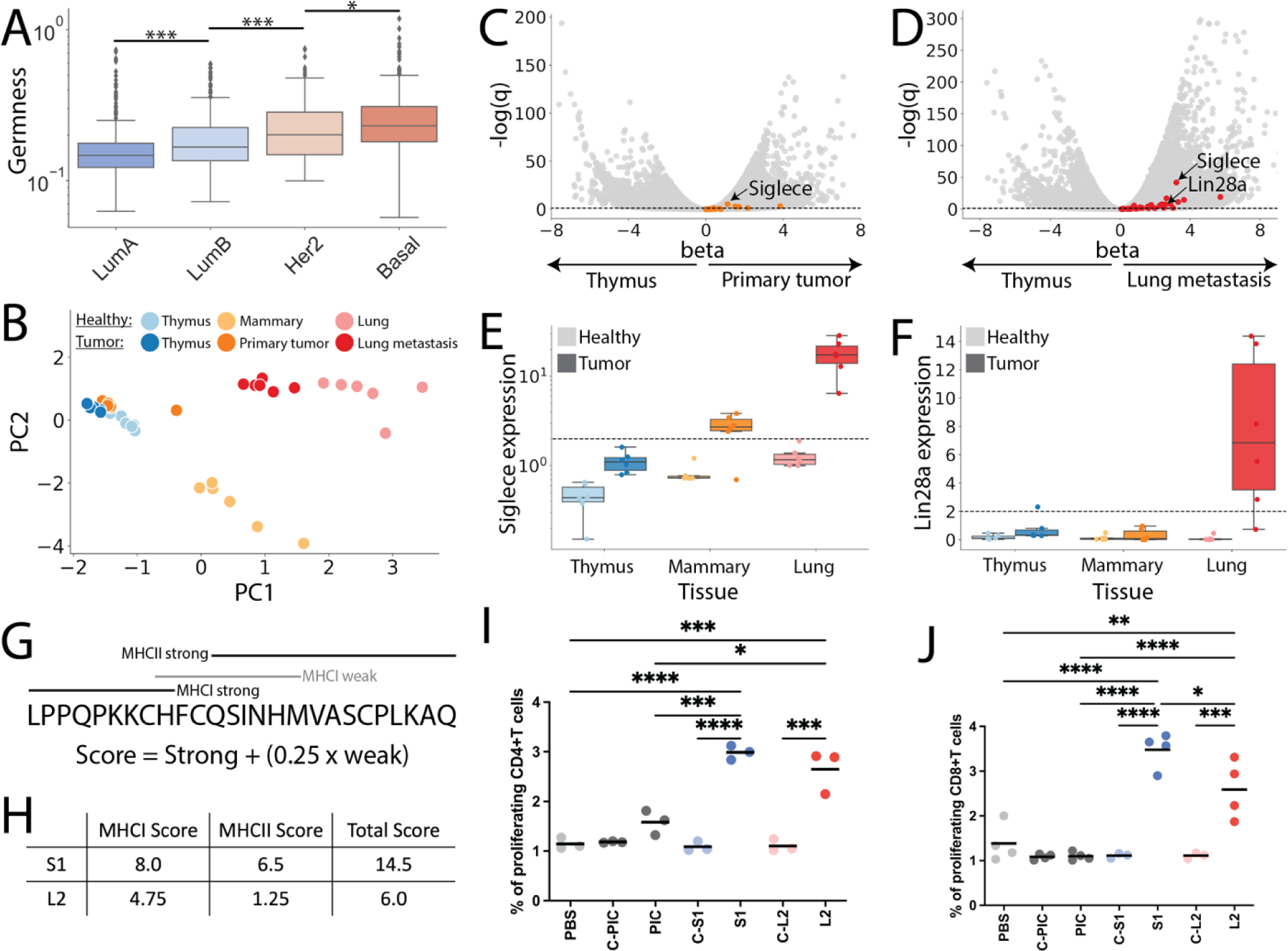
Development of CT antigen vaccines for a mouse triple-negative breast cancer model. **(A)** Germness by PAM50 subtype^56^ across GDC human breast cancer samples. The highest germness scores were observed for the basal subtype, which includes the majority of triple-negative tumors^78^. \**p*<0.05, \*\*\**p*<0.001 by Mann-Whitney U test. **(B)** Principal component (PC) analysis of thymic, mammary, and lung samples from wild type (light colors) and 4T1 tumor inoculated (dark colors) mice. **(C)** Volcano plots showing upregulated CT genes between thymus and primary tumor samples, **(D)** as well as thymus and metastatic lung tumor samples. Dotted line represents *q*=0.05. **(E)** Expression of *Siglece* (homolog of human testis-exclusive, thymic unexpressed gene *SIGLECL1*) is limited to primary murine mammary tumors and metastatic lung samples, **(F)** while *Lin28a* (homolog of *LIN28A*) is limited to metastatic lung tumors. **(G)** Germline sequences of both *Siglece* and *Lin28a* were scored according to their predicted MHCI and MHCII binding affinities. Strong binders were defined as the top 0.5% and 1% of peptides for MHCI and MHCII respectively, while weak binders represented the top 2% and 5%. Scores were calculated as the number of strong binding peptides and ¼ times the number of weak binding peptides contained within each sliding window. **(H)** We examined 2 of the top germline sequences by score as candidate CT antigen targets and synthesized these as synthetic long peptide vaccines. **(I)** Both CT antigen vaccines induced both CD4^+^ helper and **(J)** CD8^+^ cytotoxic T cell proliferation in pre-sensitized but not naïve control (c) mice. PIC-Poly(I:C), PBS-phosphate buffer solution.

We first sequenced mammary, thymus, and lung tissue samples from healthy Balb/c mice in addition to thymic, primary mammary tumor, and metastatic lung tumor samples from mice orthotopically implanted with 4T1 mammary tumor cells into the 4^th^ fat pad (**Figure 3B**). We found significant upregulation of many of our human testis-exclusive genes in both mouse primary mammary and metastatic lung tumor samples relative to paired thymic samples (**Figure 3C-D** and **Supplemental Table 4&5**). In order to identify mouse CT genes with the highest likelihood of containing immunogenic sequences, we searched for upregulated genes that were expressed in tumor samples but not in our healthy murine tissue samples, again using 2 TPM as our expression threshold (**Supplemental Figure 4** and **Supplemental Table 6**). Using these criteria, we identified two murine CT gene targets - *Siglece*, an ortholog of SIGLECL1 with expression limited to both primary mammary and metastatic lung samples, and *Lin28a*, an ortholog of LIN28A with expression observed only in lung metastases (**Figure 3E-F**).

To identify sequences to test as cancer vaccines for these two CT genes, we screened all 25-30 amino acid long substrings contained within the germline sequence of both *Siglece* and *Lin28a*. As both MHCI and MHCII epitopes significantly contribute to the anti-tumor response^57, 58^, we identified all predicted MHCI and MHCII binding epitopes contained within each of these possible sequences using NetMHCpan^37^. We then scored each germline substring by the number of predicted MHCI and MHCII epitopes, empirically weighting strong binding (top 0.5% and 1% of peptides for MHCI and MHCII, respectively) sequences more heavily than weak binding (top 2% and 5%, respectively) sequences (**Figure 3G-H**). We selected the two highest scoring distinct germline sequences for both *Siglece* (S1, S2) and *Lin28a* (L1, L2) for synthesis as peptide vaccines, ensuring that these sequences contained both epitopes that would be presented by MHCI and MHCII with hopes of stimulating robust CD4^+^ helper T cell and CD8^+^ cytotoxic T cell responses (see Methods). However, as S2 was unable to be synthesized due to hydrophobicity and L1 failed to demonstrate efficacy *in vivo* (**Supplemental Figure 5**), we focused subsequent analyses on the S1 and L2 peptide vaccines.

To confirm that these peptides can elicit T cell proliferation *in vitro*, mice were inoculated with 100,000 4T1 cells on day 0 along with pre-sensitizing subcutaneously (s.c.) with 100 µg of one of the CT peptides (S1, L2) and 100 µg of polyinosinic:polycytidylic acid (Poly(I:C) or PIC) as an adjuvant on days −4, 2, and 7. Mice injected with only Poly(I:C) or phosphate buffered saline (PBS), as well as naïve mice, were used as controls and all mice were euthanized on day 14 post-inoculation. Splenocytes were subsequently labeled with carboxyfluorescein succinimidyl ester (CSFE) and plated with PBS, PIC, S1, and L2. On day 5 after plating, proliferation indices (PI) were calculated for CD4^+^ and CD8^+^ T cells (**Figure 3I-J** and **Supplemental Figure 6**). These results demonstrated that mice pre-sensitized to both CT gene peptide sequences were able to induce robust proliferation of both CD4^+^ and CD8^+^ T cells when compared to PIC and PBS pre-treated mice, as well as naïve mice. Together, these results confirm the ability of our bioinformatic pipeline to both identify CT genes containing germline sequences that are recognized as foreign neoantigens and are capable of robustly inducing both CD4^+^ and CD8^+^ T cell responses.

### In vivo anti-tumor immune response to CTA vaccination

Given that CTA vaccination was able to induce both CD4^+^ and CD8^+^ T cell proliferation *in vitro*, we next explored the ability of the vaccines to slow primary tumor growth *in vivo*. Towards this, we investigated two different vaccination schedules, one therapeutic dosing strategy meant to mimic a treatment beginning post-diagnosis and one preventative dosing strategy that begins with a prophylactic dose. Using the therapeutic dosing strategy, we injected 100,000 4T1 cells on day 0 followed by subcutaneous injection of one of our peptide vaccines with a PIC adjuvant on days 3, 10, and 17 (**Figure 4A**). We measured the size of the primary mammary tumor following sacrifice on day 28, finding significantly smaller tumors in those mice treated with the S1, but not L2, peptide vaccine (**Figure 4B**). Using the preventative dosing strategy with vaccination occurring on days −4, 2, 7, and 14, we again found significantly smaller primary tumors in mice treated with the S1 peptide (**Figure 4C-D**). Mice treated with S1 using the preventative dosing strategy tended to have smaller primary tumors at day 28 when compared to therapeutic dosing (403 ±148mg vs 604 ± 272mg [mean ± standard deviation], p=0.18 by Mann-Whitney U test). Of note, we did not expect primary mammary tumor growth to be altered by the L2 vaccine given the lack of *Lin28a* expression in primary tumors (**Figure 3E-F**). These findings directly demonstrate the ability of S1 vaccination to slow *in vivo* tumor growth in an expression-dependent fashion.

**Figure 4.**
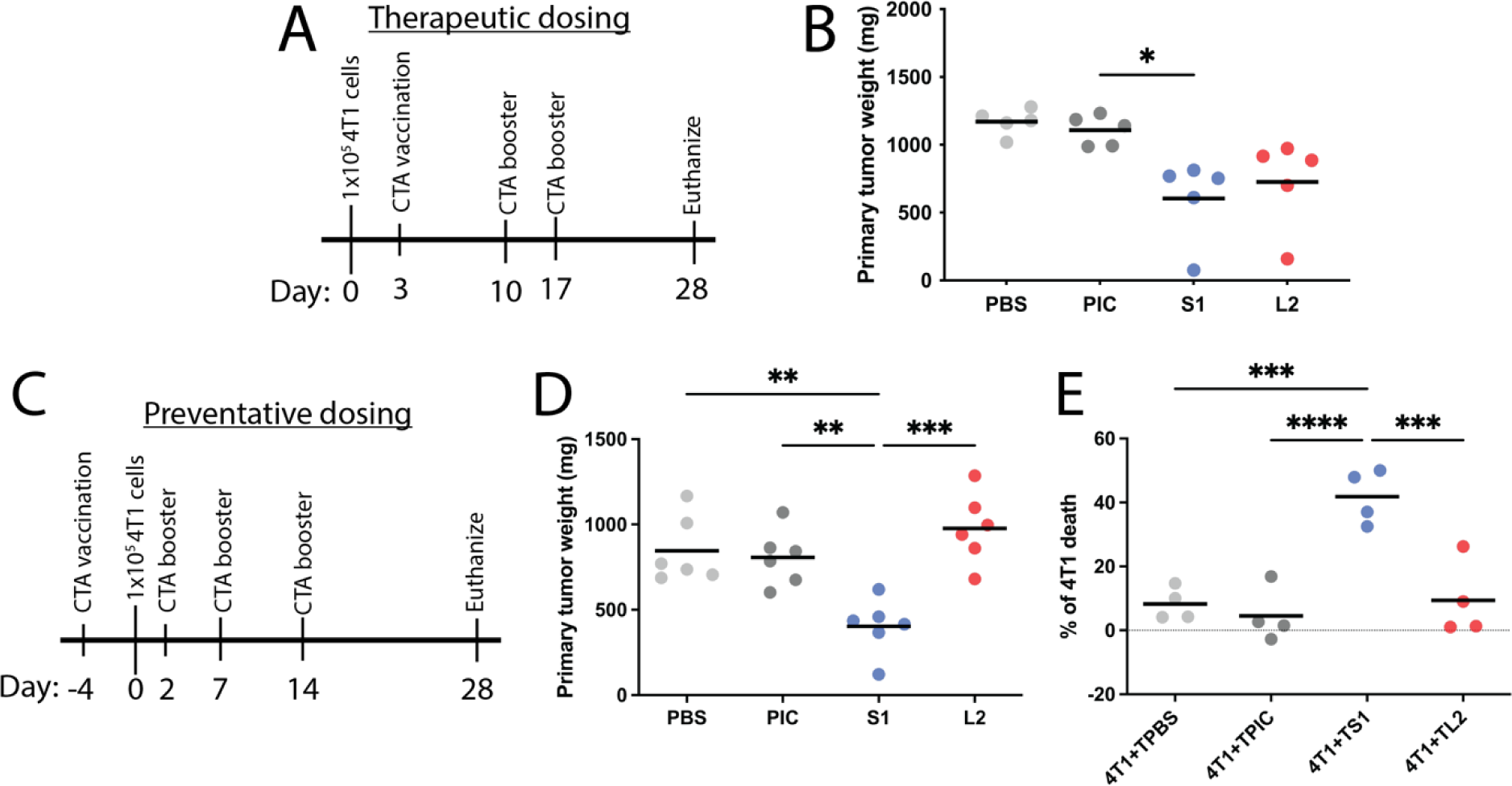
CT antigen vaccination reduces primary tumor growth by inducing targeted 4T1 cell death. **(A)** Schematic demonstrating therapeutic treatment dosing strategy with vaccination starting on day 3 following 4T1 tumor inoculation. Additional CT antigen vaccine boosters were administered on days 10 and 17 prior to euthanization on day 28. **(B)** Primary mammary tumor weights following sacrifice on day 28 following treatment with Siglece (S) and Lin28a (L) CT antigen vaccines following the therapeutic dosing regimen. **(C)** Alternatively, the prevention vaccination regimen had initial CT antigen vaccination 4 days prior to tumor inoculation, with subsequent boosters provided on days 2, 7, and 14 prior to euthanization again on day 28. **(D)** Mice treated with S1, but not L2, CT antigen vaccine according to the preventative dosing strategy had significantly lower primary tumor size relative to Poly(I:C) and phosphate buffer solution (PBS) controls. **(E)** 4T1 cell death was significantly increased when co-cultured with tumor infiltrates from mice treated with the S1 vaccine according to the preventative dosing strategy. No increase in cell death was observed with tumor infiltrates from mice treated with L2, consistent with Siglece but not Lin28a expression in primary mammary tumors. For all panels, *p<0.05, **p<0.01, ***p<0.001 by one-way ANOVA through Tukey’s *post hoc* tests, all other pairwise comparisons not statistically significant.

To confirm that CT antigen vaccination slows primary tumor growth by increasing immune recognition and subsequent cytolysis of tumor cells, we next performed a functional cytotoxic assay using tumor infiltrates. We co-cultured CSFE-labeled primary 4T1 cells for 24 hours with tumor cells isolated from primary mammary tumors treated with the prevention dosing strategy in a ratio of 5:1 of tumor cells to 4T1 cells. We then used flow cytometry to quantify the percentage of cell death among the population of CSFE-labeled primary 4T1 cells, adjusting for the basal level of cell death of 4T1 cells cultured alone. We found significantly elevated rates of 4T1 cell death when exposed to tumor cells of mice treated with S1 (**Figure 4E** and **Supplemental Figure 7**). As expected, given the lack of *Lin28a* expression in primary tumors, no increase in primary 4T1 cell death was observed when exposed to infiltrates treated with the *Lin28a* vaccine (**Figure 4E**).

Given the cytotoxic abilities of S1-treated tumor infiltrates, we next explored T cell migration and function in the tumor immune microenvironment on day 28 following the preventative dosing schedule. We first used flow cytometry to quantify the percentage of cytotoxic CD8^+^ T cells within tumor cell isolates relative to the total number of CD45^+^ leukocytes. While S1 vaccination tended to have higher relative fractions of cytotoxic T cells, this effect was not significant. Interestingly, L2 vaccination showed a significant increase in the proportion of CD8^+^ T cells relative to PIC and PBS treated controls (**Figure 5A**). We additionally found that vaccination with both S1 and L2 increased the frequency of CD4^+^ T cells among all CD45^+^ leukocytes relative to PBS alone (**Figure 5B**).

**Figure 5.**
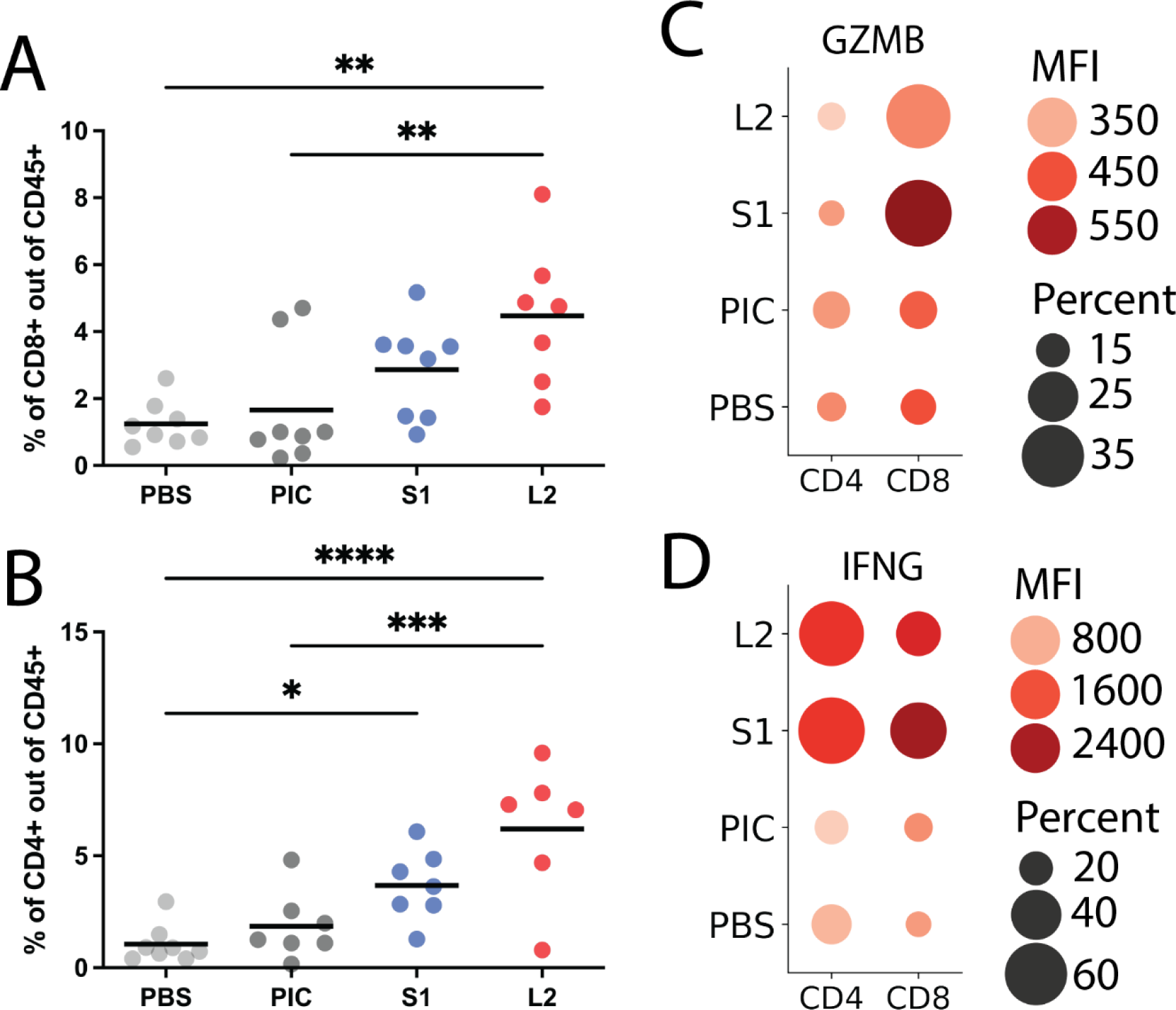
CT antigen vaccination stimulates a robust CD4^+^ and CD8^+^ T cell anti-tumor immune response. **(A)** Tumors treated with the L2 vaccine according to the prevention dosing regimen had a significantly increased proportion of CD8^+^ cytotoxic T cells among tumor infiltrating CD45^+^ lymphocytes. **(B)** Similarly, a higher proportion of tumor infiltrating CD4^+^ helper T cells relative to all CD45^+^ lymphocytes was observed following S1 and L2 preventative dosing. **(C)** Dot plot demonstrating a significantly higher percentage of granzyme B producing CD8^+^ T cells following treatment with either the S1 or L2 vaccines found in tumors. Increasing dot size is correlated with an increasing percentage of cells expressing granzyme B while dot color corresponds to the average expression level as indicated by mean fluorescent intensity (MFI). **(D)** Similarly, a significantly higher fraction of CD4+ and CD8+ T cells expressed interferon gamma following both S1 and L2 vaccination, with significantly higher average expression seen in CD4^+^ T cells for both vaccines. For all panels, *p<0.05, **p<0.01, ***p<0.001 by one-way ANOVA through Tukey’s *post hoc* tests, all other pairwise comparisons not statistically significant.

To examine the functionality of these tumor-infiltrating lymphocytes, we additionally compared expression of granzyme B (GZMB) and interferon-gamma (IFNG) by treatment group. Primary mammary tumors in mice treated with both S1 and L2 had significantly higher percentage of GZMB producing CD8^+^ T cells within the tumor microenvironment. However, vaccination did not significantly increase the average expression level of GZMB amongst tumor-infiltrating CD8^+^ T cells (**Figure 5C** and **Supplemental Figure 8**). In contrast, S1 and L2 vaccination significantly increased the proportion of IFNG producing CD4^+^ T cells, as well as increased the average expression of IFNG within these CD4^+^ T cells (**Figure 5D** and **Supplemental Figure 9**). No differences were observed in the proportion of cells expressing or in the average expression level of GZMB and IFNG between mice vaccinated with S1 and L2. In summary, these findings demonstrate that the S1 vaccine induces robust CD4^+^ and CD8^+^ T cell anti-tumor functions that contributes to increased tumor cell death and reduced tumor growth in this mouse model of TNBC.

### CTA vaccination drastically reduces the number of pulmonary metastases

Both *Siglece* and *Lin28a* have significant expression in lung metastases in this 4T1 mouse model of TNBC (**Figure 3E-F**). We therefore asked whether CT gene vaccination could be used to slow or prevent the development of metastatic disease. Gross pathologic examination of lungs harvested from day 28 mice treated with the preventative dosing vaccination schedule demonstrated a significant reduction in metastatic disease burden in mice treated with either S1 or L2 (**Figure 6A**). This reduction in the number of lung metastases was further supported by histologic analysis (**Figure 6B**). Quantification of the number of observable lung metastases in each sample confirmed a significant reduction in the number of metastatic sites in those mice treated with either the S1 or L2 peptide vaccine (**Figure 6C**).

**Figure 6.**
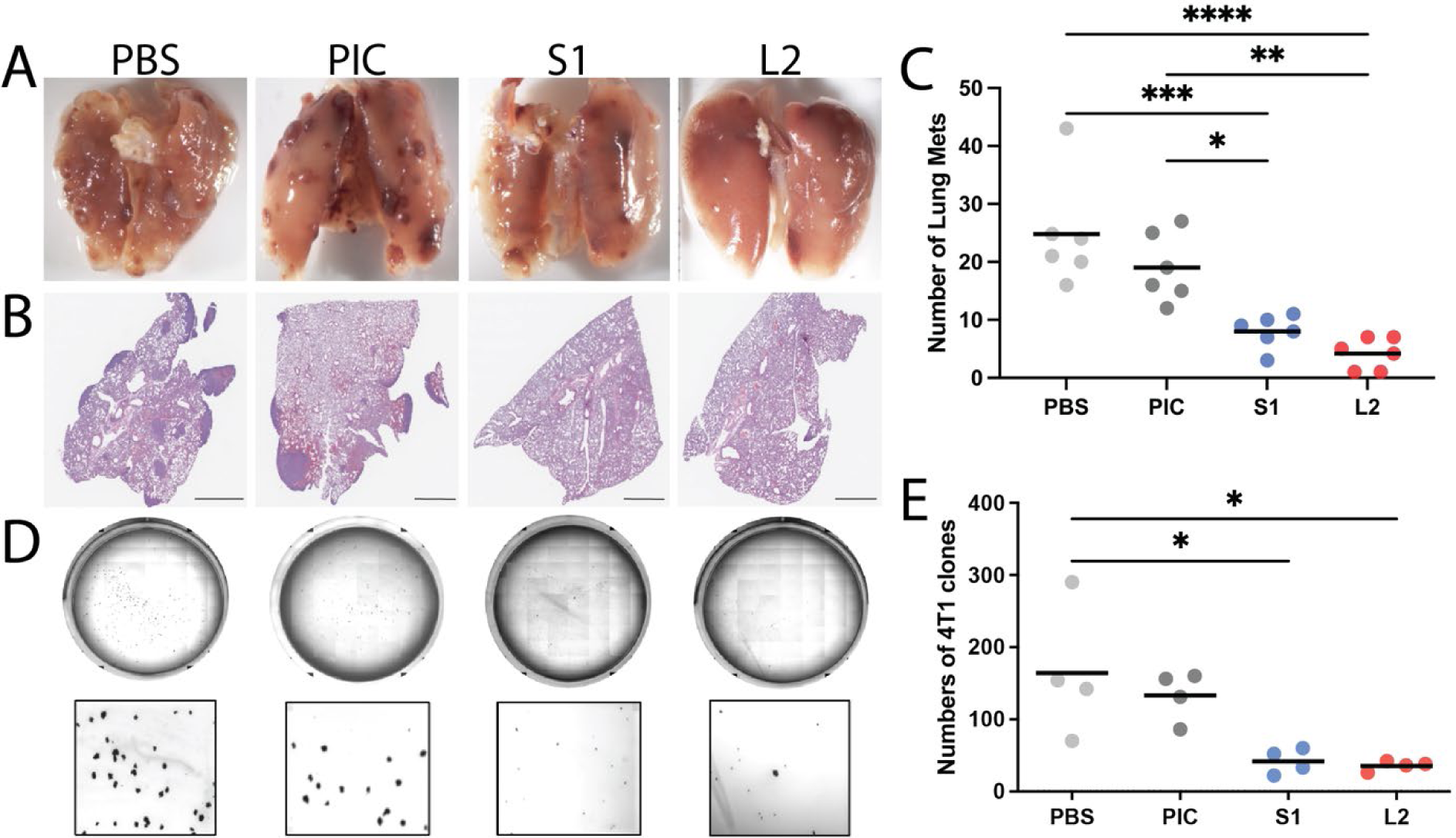
CT antigen vaccination significantly reduces lung metastases. **(A)** Representative photos showing the extent of lung metastases on gross examination and **(B)** histological examination with hematoxylin and eosin (H&E) staining following harvest on day 28 in mice treated following the preventative dosing strategy. Scale bars correspond to 2mm. **(C)** Quantification demonstrates significantly reduced numbers of lung metastases following treatment with either S1 or L2 CT antigen vaccines. **(D)** Representative photos demonstrating the number of 4T1 colonies formed from lung tissue harvested on day 28 following the preventative dosing strategy. **(E)** The number of 4T1 colonies was significantly reduced following treatment with the S1 and L2 vaccines. For all panels, *p<0.05, **p<0.01, ***p<0.001 by one-way ANOVA through Tukey’s *post hoc* tests, all other pairwise comparisons not statistically significant.

To further characterize the ability of CT antigen vaccination to reduce the number of lung metastases, we performed a clonogenic assay. Briefly, mice were again treated according to the prevention strategy and single-cell suspensions made from lungs harvested on day 28. We then plated 200 cells in complete media supplemented with 6-thioguanine and subsequently counted the number of 4T1 colonies after 14 days (**Figure 6D**). Quantification of the number of colonies from each treatment type again demonstrated a significant reduction in the numbers of lung 4T1 colonies from those mice treated with either the S1 or L2 peptide vaccine (**Figure 6E**). These findings are again indicative of a lower overall metastatic disease burden in mice treated with our CT antigen vaccines. Altogether, findings show a striking and significant reduction in the number of lung metastases following vaccination.

## Conclusions

The capacity of the immune system to specifically attack neoplastic cells renders it a powerful weapon for the long-term control of many different cancer types. T cell recognition of high-quality neoantigens underlies natural anti-tumor immunity and response to immunotherapy^59–61^. Current immune checkpoint blockade strategies have transformed our understanding of the tumor immune microenvironment and the clinical treatment of many high tumor mutational burden cancer types. However, these immunotherapies do not allow for targeted treatment of specific tumor-specific neoantigens, with clinical use consequently often limited by off-target autoimmunity^62^. Personalized cancer vaccines, which allow for induction of neoantigen-specific immunity, have shown promise in early clinical trials^3, 4^. These somatic mutation neoantigen targets, however, are frequently unique to each tumor and therefore clinical treatment requires labor-intensive bioinformatic analyses to design and subsequently synthesize a custom vaccine for each patient. The ability to target immunogenic germline sequences would therefore enable the design of common vaccines applicable to all tumors expressing that gene, with CT genes representing a particularly intriguing source. Indeed, cancer vaccines targeting CTA have shown promise in melanoma^63–66^, esophageal^67^, gastric^68, 69^, head and neck^70^, biliary^71, 72^, and pancreatic^73^ cancers, though recent phase 3 trials targeting melanoma-associated antigen 3 (MAGE-A3) in melanoma^74^ and NSCLC^75^ failed to meet their primary end points.

Despite this interest in the use of CT genes as neoantigen targets for immunotherapy^14–16^, there has been no consensus method for identifying CT genes^29–31^. The GTEx and TCGA sequencing repositories have enabled high-throughput screening for CT gene expression patterns^35, 36^, though these studies have not focused on the identification of immunogenic targets for immunotherapy. In this study, we used the Phi Correlation Coefficient^41^ with a highly conservative threshold to identify genes with testis-exclusive expression across the GTEx healthy tissue database. We additionally screened this set of testis-exclusive genes for human thymic expression^43^, further narrowing our set of genes without detectable thymic expression and consequently the most likely to lack central tolerance. Consequently, we defined our final set of 103 CT genes as those testis-exclusive genes without thymic expression that were expressed in >5% of tumor samples in at least 2 separate tumor types. Together, this bioinformatic pipeline aimed to identify a set of CT genes that were likely to serve as both immunogenic and generalizable antigen targets. We then confirmed that expression of these CT genes in the GDC pan-cancer cohort was correlated with more advanced tumor stage and markers of anti-tumor immunity, as well as overall survival.

TNBC is extremely aggressive and associated with poor prognosis and higher risk of early recurrence and metastasis. Because TNBC lacks targetable hormone receptor expression, the standard-of-care is chemotherapy. Yet, 50% of TNBC patients are insensitive to chemotherapy and most patients relapse and develop metastases within 3 years. Thus, a comprehensive analysis using a larger cohort of samples from various stages would be the necessary next step. Nonetheless, in support of these findings, we developed peptide vaccines against *Siglece* and *Lin28a* that are expressed in the 4T1 mouse model of TNBC. Mice sensitized with a peptide vaccine targeting Siglece had significantly reduced primary tumor sizes. We further demonstrated that this reduction in primary tumor growth was mediated by both CD4^+^ and CD8^+^ T cell function and subsequent tumor cell death. We further found that vaccination against both Siglece and Lin28a, which are expressed by metastatic tumors, significantly reduced the number of lung metastases in mice. Siglece and Lin28a are newly identified CT antigens in cancer. Although Lin28a/b have been previously implicated in breast cancer metastasis^76, 77^, little is known of Siglece function.

CTAs are normally expressed in the testis but are highly expressed across cancer and associated with disease stage, an unfavorable prognosis, and cancer invasion, making them potentially promising targets. Quite strikingly, CTA vaccination revealed induction of both CD4+ and CD8+ T cell responses, which is a promising direction for the future development of peptide vaccines for immunotherapy. Additional studies are needed to explore new CTAs with high immunogenicity and specificity that can serve as ideal targets for cancer immunotherapy^16^.

In summary, CT genes represent an intriguing source of targetable antigens that may enable the development of new immunotherapeutics. In this study, we implement a novel bioinformatic pipeline that focuses on the high-throughput screening of the GTEx and TCGA databases to identify high-confidence testis-exclusive genes that are likely to lack central tolerance and are robustly expressed across at least two tumor types. We validate the immunogenicity of two CT genes using a mouse model of TNBC, with vaccination inducing tumor-specific immunity in an expression-dependent manner. Further exploration of these immunogenic CT genes as vaccination targets in human tumors will be of particular interest, in addition to the further identification and validation of therapeutic antigen-based vaccines that target immunogenic tumor-specific CT genes for the treatment of TNBC.

## Acknowledgements

JAC was partially supported by NIHGM MSTP Training award T32-GM008444. GA was funded by the Simons Foundation, a LIBH grant, and the Stand Up To Cancer-Breast Cancer Research Foundation Convergence Team Translational Grant Number 310 SU2C-BCRF 2015-001. BJB was funded by the NCI 1R21CA195256-01, Department of Defense Breast Cancer Research Program W81XWH-19-1-0113, the Manhasset Women’s Coalition Against Breast Cancer and a Northwell-Cold Spring Harbor Collaborative Research Grant.

## References

1. Tran, E., Robbins, P. F. & Rosenberg, S. A. Final common pathway’ of human cancer immunotherapy: Targeting random somatic mutations. Nat Immunol 18, 255–262 (2017).

2. Schumacher, T. N. & Schreiber, R. D. Neoantigens in cancer immunotherapy. Science (1979) 348, 69–74 (2015).

3. Saxena, M., van der Burg, S. H., Melief, C. J. M. & Bhardwaj, N. Therapeutic cancer vaccines. Nat Rev Cancer 21, 360–378 (2021).

4. Blass, E. & Ott, P. A. Advances in the development of personalized neoantigen-based therapeutic cancer vaccines. Nat Rev Clin Oncol 18, 215–229 (2021).

5. Ott, P. A. et al. A Phase Ib Trial of Personalized Neoantigen Therapy Plus Anti-PD-1 in Patients with Advanced Melanoma, Non-small Cell Lung Cancer, or Bladder Cancer. Cell 183, 347–362.e24 (2020).

6. Ott, P. A. et al. An immunogenic personal neoantigen vaccine for patients with melanoma. Nature 547, 217–221 (2017).

7. Sahin, U. et al. Personalized RNA mutanome vaccines mobilize poly-specific therapeutic immunity against cancer. Nature 547, 222–226 (2017).

8. Carreno, B. M. et al. Cancer immunotherapy. A dendritic cell vaccine increases the breadth and diversity of melanoma neoantigen-specific T cells. Science 348, 803–808 (2015).

9. Hilf, N. et al. Actively personalized vaccination trial for newly diagnosed glioblastoma. Nature 565, 240–245 (2019).

10. Keskin, D. B. et al. Neoantigen vaccine generates intratumoral T cell responses in phase Ib glioblastoma trial. Nature 565, 234–239 (2019).

11. Schumacher, T. et al. A vaccine targeting mutant IDH1 induces antitumour immunity. Nature 512, 324–327 (2014).

12. Ozlem, T. et al. Targeting the Heterogeneity of Cancer with Individualized Neoepitope Vaccines. Clin Cancer Res 22, 1885–1896 (2016).

13. Pearlman, A. H. et al. Targeting public neoantigens for cancer immunotherapy. Nat Cancer 2, 487–497 (2021).

14. Gjerstorff, M. F., Burns, J. & Ditzel, H. J. Cancergermline antigen vaccines and epigenetic enhancers: Future strategies for cancer treatment. Expert Opin Biol Ther 10, 1061–1075 (2010).

15. Gjerstorff, M. F., Andersen, M. H. & Ditzel, H. J. Oncogenic cancer/testis antigens: prime candidates for immunotherapy. Oncotarget 6, 1577215787 (2015).

16. Wei, X. et al. Cancer-Testis Antigen Peptide Vaccine for Cancer Immunotherapy: Progress and Prospects. Transl Oncol 12, 733–738 (2019).

17. Scanlan, M. J., Gure, A. O., Jungbluth, A. A., Old, L. J. & Chen, Y. T. Cancer/testis antigens: an expanding family of targets for cancer immunotherapy. Immunol Rev 188, 22–32 (2002).

18. Batra, R. N. et al. DNA methylation landscapes of 1538 breast cancers reveal a replication-linked clock, epigenomic instability and cis-regulation. Nat Commun 12, (2021).

19. Maxfield, K. E. et al. Comprehensive functional characterization of cancer-testis antigens defines obligate participation in multiple hallmarks of cancer. Nat Commun 6, (2015).

20. Suyama, T. et al. Expression of Cancer/Testis Antigens in Prostate Cancer is Associated With Disease Progression. Prostate 70, 1778 (2010).

21. Barrow, C. et al. Tumor antigen expression in melanoma varies according to antigen and stage. Clin Cancer Res 12, 764–771 (2006).

22. Rousseaux, S. et al. Ectopic activation of germline and placental genes identifies aggressive metastasis-prone lung cancers. Sci Transl Med 5, 186ra66 (2013).

23. Shukla, S. A. et al. Cancer-Germline Antigen Expression Discriminates Clinical Outcome to CTLA−4 Blockade. Cell 173, 624–633.e8 (2018).

24. Yao, J. et al. Tumor subtype-specific cancer-testis antigens as potential biomarkers and immunotherapeutic targets for cancers. Cancer Immunol Res 2, 371–379 (2014).

25. Mahmoud, A. M. Cancer testis antigens as immunogenic and oncogenic targets in breast cancer. Immunotherapy 10, 769–778 (2018).

26. Lam, R. A. et al. Cancer-testis antigens in triple-negative breast cancer: Role and potential utility in clinical practice. Cancers (Basel) 13, 3875 (2021).

27. Chen, Y. T. et al. Multiple cancer/testis antigens are preferentially expressed in hormone-receptor negative and high-grade breast cancers. PLoS One 6, (2011).

28. Almeida, L. G. et al. CTdatabase: a knowledge-base of high-throughput and curated data on cancer-testis antigens. Nucleic Acids Res 37, D816–D819 (2009).

29. Hofmann, O. et al. Genome-wide analysis of cancer/testis gene expression. PNAS 105, 20422– 20427 (2008).

30. Chen, Y.-T. et al. Identification of cancer/testis-antigen genes by massively parallel signature sequencing. PNAS 102, 7940–7945 (2005).

31. Taguchi, A. et al. A Search for Novel Cancer/Testis Antigens in Lung Cancer Identifies VCX/Y Genes, Expanding the Repertoire of Potential Immunotherapeutic Targets. Cancer Res 74, 4694–4705 (2014).

32. Lonsdale, J. et al. The Genotype-Tissue Expression (GTEx) project. Nature Genetics 2013 45:6 45, 580–585 (2013).

33. Weinstein, J. N. et al. The Cancer Genome Atlas Pan-Cancer analysis project. Nature Genetics 2013 45:10 45, 1113–1120 (2013).

34. Zhang, Z. et al. Uniform genomic data analysis in the NCI Genomic Data Commons. Nat Commun 12, 1226 (2021).

35. Wang, C. et al. Systematic identification of genes with a cancer-testis expression pattern in 19 cancer types. Nat Commun 7, (2016).

36. Lira Da Silva, V., et al. Genome-wide identification of cancer/testis genes and their association with prognosis in a pan-cancer analysis. Oncotarget 8, 92966–92977 (2017).

37. Reynisson, B., Alvarez, B., Paul, S., Peters, B. & Nielsen, M. NetMHCpan−4.1 and NetMHCIIpan−4.0: improved predictions of MHC antigen presentation by concurrent motif deconvolution and integration of MS MHC eluted ligand data. Nucleic Acids Res 48, W449–W454 (2020).

38. Desrichard, A., Snyder, A. & Chan, T. A. Cancer Neoantigens and Applications for Immunotherapy. Clin Cancer Res 22, 807–812 (2016).

39. Cunningham, F. et al. Ensembl 2022. Nucleic Acids Res 50, D988–D995 (2022).

40. Wagner, G. P., Kin, K. & Lynch, V. J. A model based criterion for gene expression calls using RNA-seq data. Theory Biosci 132, 159–164 (2013).

41. Melé, M. et al. The human transcriptome across tissues and individuals. Science (1979) 348, 660–665 (2015).

42. Mi, H. et al. Protocol Update for large-scale genome and gene function analysis with the PANTHER classification system (v.14.0). Nat Protoc 14, 703–721 (2019).

43. Carter, J. A. et al. Transcriptomic diversity in human medullary thymic epithelial cells. Nature Communications 2022 13:1 13, 1–15 (2022).

44. Coles, A. J., et al. Keratinocyte growth factor impairs human thymic recovery from lymphopenia. JCI Insight 4, (2019).

45. Bray, N. L., Pimentel, H., Melsted, P. & Pachter, L. Near-optimal probabilistic RNA-seq quantification. Nat Biotechnol 34, 525–527 (2016).

46. Grossman, R. L. et al. Toward a Shared Vision for Cancer Genomic Data. New England Journal of Medicine 375, 1109–1112 (2016).

47. Thorsson, V. et al. The Immune Landscape of Cancer. Immunity 48, 812–830.e14 (2018).

48. Davidson-Pilon, C. lifelines: survival analysis in Python. J Open Source Softw 4, 1317 (2019).

49. Pimentel, H., Bray, N. L., Puente, S., Melsted, P. & Pachter, L. Differential analysis of RNA-seq incorporating quantification uncertainty. Nat Methods 14, 687–690 (2017).

50. Mi, H. et al. Protocol Update for large-scale genome and gene function analysis with the PANTHER classification system (v.14.0). Nature Protocols 2019 14:3 14, 703–721 (2019).

51. Thomas, P. D. et al. PANTHER: Making genome-scale phylogenetics accessible to all. Protein Science 31, 8–22 (2022).

52. Thorsson, V. et al. The Immune Landscape of Cancer. Immunity 48, 812–830.e14 (2018).

53. Yang, P., Huo, Z., Liao, H. & Zhou, Q. Cancer/testis antigens trigger epithelial-mesenchymal transition and genesis of cancer stem-like cells. Curr Pharm Des 21, 1292–1300 (2015).

54. Chen, Y. T. et al. Multiple Cancer/Testis Antigens Are Preferentially Expressed in Hormone-Receptor Negative and High-Grade Breast Cancers. PLoS One 6, e17876 (2011).

55. Bernard, P. S. et al. Supervised risk predictor of breast cancer based on intrinsic subtypes. Journal of Clinical Oncology 27, 1160–1167 (2009).

56. Koboldt, D. C. et al. Comprehensive molecular portraits of human breast tumours. Nature 2012 490:7418 490, 61–70 (2012).

57. Tran, E. et al. Cancer immunotherapy based on mutation-specific CD4+ T cells in a patient with epithelial cancer. Science (1979) 344, 641–645 (2014).

58. Kreiter, S. et al. Mutant MHC class II epitopes drive therapeutic immune responses to cancer. Nature 2015 520:7549 520, 692–696 (2015).

59. Balachandran, V. P. et al. Identification of unique neoantigen qualities in long-term survivors of pancreatic cancer. Nature 2017 551:7681 551, 512–516 (2017).

60. Łuksza, M. et al. Neoantigen quality predicts immunoediting in survivors of pancreatic cancer. Nature 2022 606:7913 606, 389–395 (2022).

61. Luksza, M. et al. A neoantigen fitness model predicts tumour response to checkpoint blockade immunotherapy. Nature 2017 551:7681 551, 517–520 (2017).

62. Hodi, F. S. et al. Nivolumab plus ipilimumab or nivolumab alone versus ipilimumab alone in advanced melanoma (CheckMate 067): 4-year outcomes of a multicentre, randomised, phase 3 trial. Lancet Oncol 19, 1480–1492 (2018).

63. Lattanzi, M. et al. Adjuvant NY-ESO-1 vaccine immunotherapy in high-risk resected melanoma: A retrospective cohort analysis. J Immunother Cancer 6, 1–10 (2018).

64. Marchand, M. et al. Tumor regression responses in melanoma patients treated with a peptide encoded by gene MAGE-3. Int J Cancer 63, 883–885 (1995).

65. Kruit, W. H. J. et al. Phase 1/2 study of subcutaneous and intradermal immunization with a recombinant MAGE-3 protein in patients with detectable metastatic melanoma. Int J Cancer 117, 596–604 (2005).

66. McQuade, J. L. et al. A phase II trial of recombinant MAGE-A3 protein with immunostimulant AS15 in combination with high-dose Interleukin-2 (HDIL2) induction therapy in metastatic melanoma. BMC Cancer 18, (2018).

67. Kono, K. et al. Multicenter, phase II clinical trial of cancer vaccination for advanced esophageal cancer with three peptides derived from novel cancer-testis antigens. J Transl Med 10, (2012).

68. Higashihara, Y. et al. Phase I clinical trial of peptide vaccination with URLC10 and VEGFR1 epitope peptides in patients with advanced gastric cancer. Int J Oncol 44, 662–668 (2014).

69. Ishikawa, H. et al. Phase I clinical trial of vaccination with LY6K-derived peptide in patients with advanced gastric cancer. Gastric Cancer 17, 173–180 (2014).

70. Yoshitake, Y. et al. Phase II clinical trial of multiple peptide vaccination for advanced head and neck cancer patients revealed induction of immune responses and improved OS. Clin Cancer Res 21, 312–321 (2015).

71. Aruga, A. et al. Phase I clinical trial of multiple-peptide vaccination for patients with advanced biliary tract cancer. J Transl Med 7, 61 (2014).

72. Aruga, A. et al. Long-term vaccination with multiple peptides derived from cancer-testis antigens can maintain a specific T-cell response and achieve disease stability in advanced biliary tract cancer. Clinical Cancer Research 19, 2224–2231 (2013).

73. Okuyama, R., Aruga, A., Hatori, T., Takeda, K. & Yamamoto, M. Immunological responses to a multi-peptide vaccine targeting cancer-testis antigens and VEGFRs in advanced pancreatic cancer patients. Oncoimmunology 2, (2013).

74. Dreno, B. et al. MAGE-A3 immunotherapeutic as adjuvant therapy for patients with resected, MAGE-A3-positive, stage III melanoma (DERMA): a double-blind, randomised, placebo-controlled, phase 3 trial. Lancet Oncol 19, 916–929 (2018).

75. Vansteenkiste, J. F. et al. Efficacy of the MAGE-A3 cancer immunotherapeutic as adjuvant therapy in patients with resected MAGE-A3-positive non-small-cell lung cancer (MAGRIT): a randomised, double-blind, placebo-controlled, phase 3 trial. Lancet Oncol 17, 822–835 (2016).

76. Wu, K., Ahmad, T. & Eri, R. LIN28A: A multifunctional versatile molecule with future therapeutic potential. World J Biol Chem 13, 35–46 (2022).

77. Qi, M. et al. Lin28B-high breast cancer cells promote immune suppression in the lung pre-metastatic niche via exosomes and support cancer progression. Nat Commun 13, 897 (2022).

78. Badve, S. et al. Basal-like and triple-negative breast cancers: a critical review with an emphasis on the implications for pathologists and oncologists. Modern Pathology 2011 24:2 24, 157–167 (2010).

