## Supplemental Figures for "Identification of pan-cancer/testis genes and validation of therapeutic targeting in triple-negative breast cancer: Lin28a- and Siglece-based vaccination induces anti-tumor immunity and inhibits metastasis"

**Supplemental Materials**

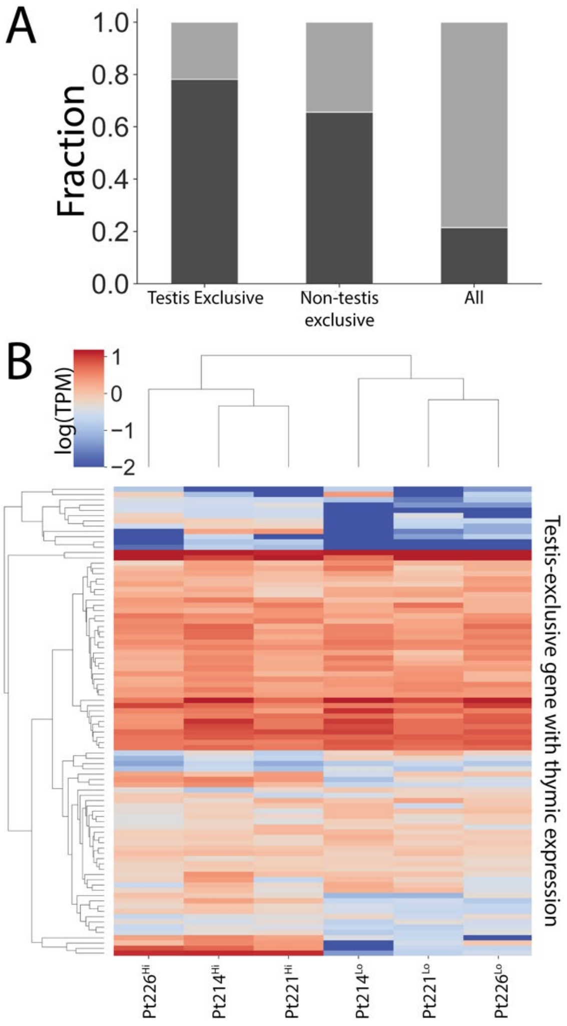


**Supplemental Figure 1. Thymic expression of testis-exclusive genes. (A)** Testis-exclusive genes were less likely to lack thymic expression relative to non-testis tissue-exclusive genes (odds ratio [OR] 1.9, $p$=*0.12* by Fischer's exact test) and all (i.e. non-tissue exclusive genes) genes (OR 13.1, $p=1.8\text{x}10^{-128}$). Non-testis tissue-exclusive genes were less frequently expressed in the thymus relative to non-tissue exclusive genes (OR 7.0, $p=1.0\text{x}10^{-7}$). **(B)** No systematic differences were observed in the expression of testis-exclusive genes with thymic expression between immature MHCII low mTECs and mature MHCII high mTECs.

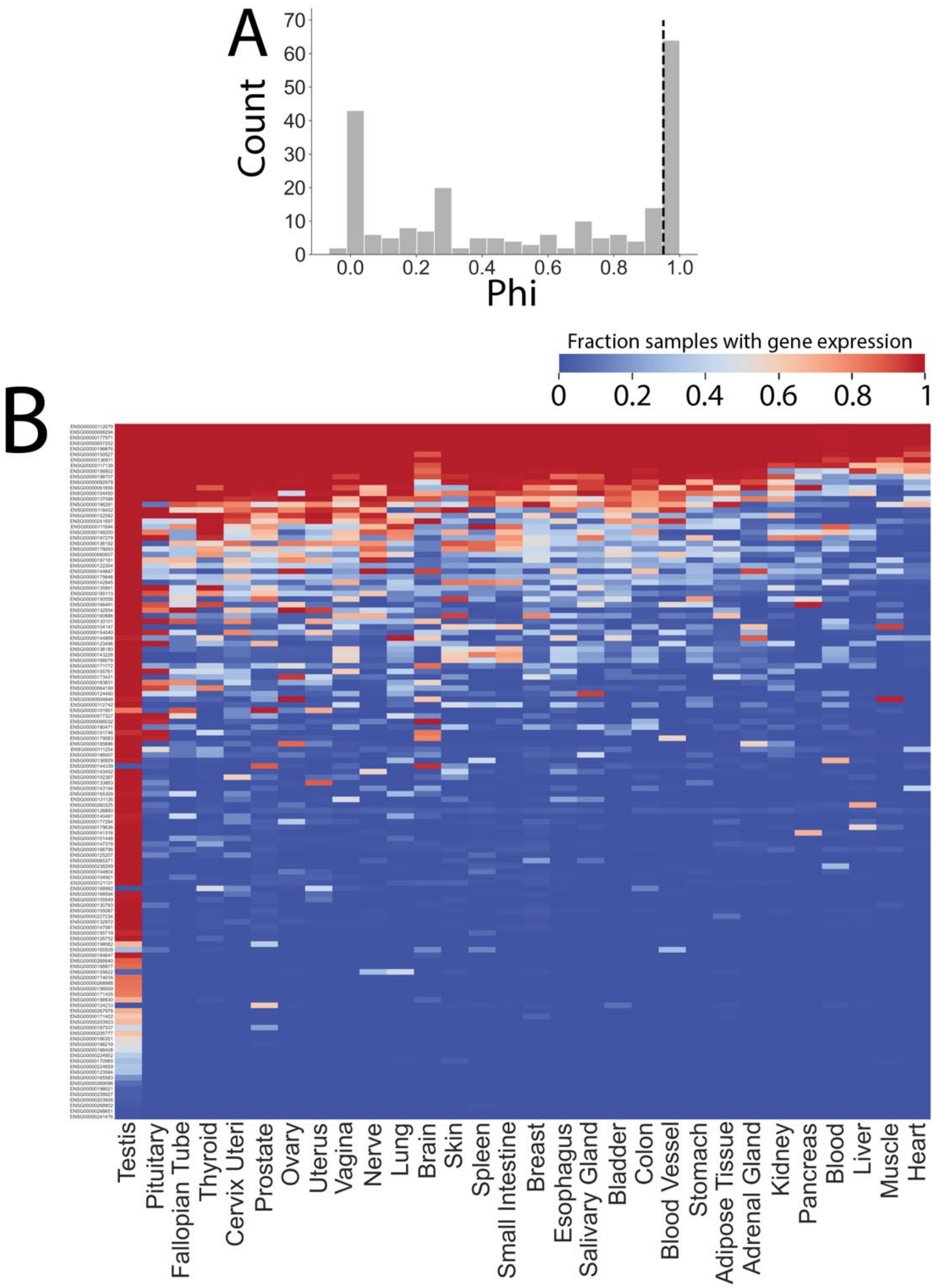


**Supplemental Figure 2. Expression of CTdatabase genes that do not meet our criteria for testis-exclusive expression. (A)** Distribution of testis Phi values for CTdatabase genes, with a threshold of 0.95 (dotted line) indicating our threshold for testis-exclusive expression. We found that 157 out of 221 (71%) of CTdatabase genes did not meet our threshold for testis-exclusive expression. **(B)** Of these 157 CTdatabase genes, 32 had no detectable expression in any tissue type. The fraction of healthy GTEx samples expressing each of the remaining 125 CTdatabase genes without tissue-exclusive expression are shown by tissue type.

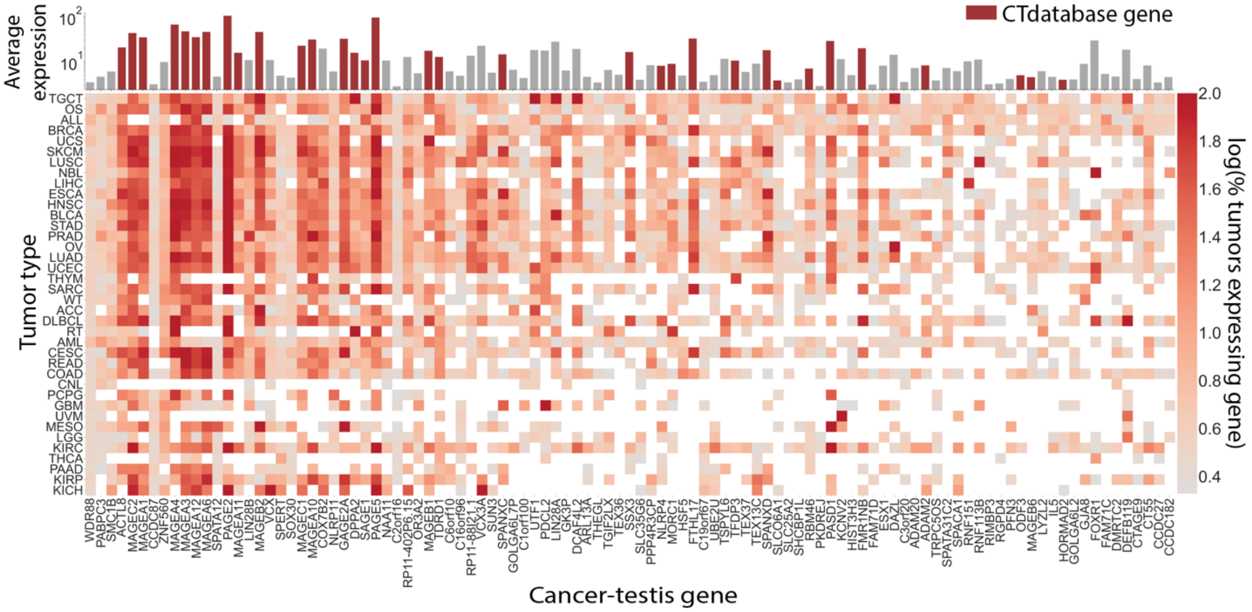


**Supplemental Figure 3. Tumor expression of cancer-testis genes.** The average expression, as transcripts per million, for each of the 103 cancer-testis genes is shown across GDC tumor types. The average expression across all cancer types is shown in the top bar graph, with those genes present in the CTdatabase shown in red.

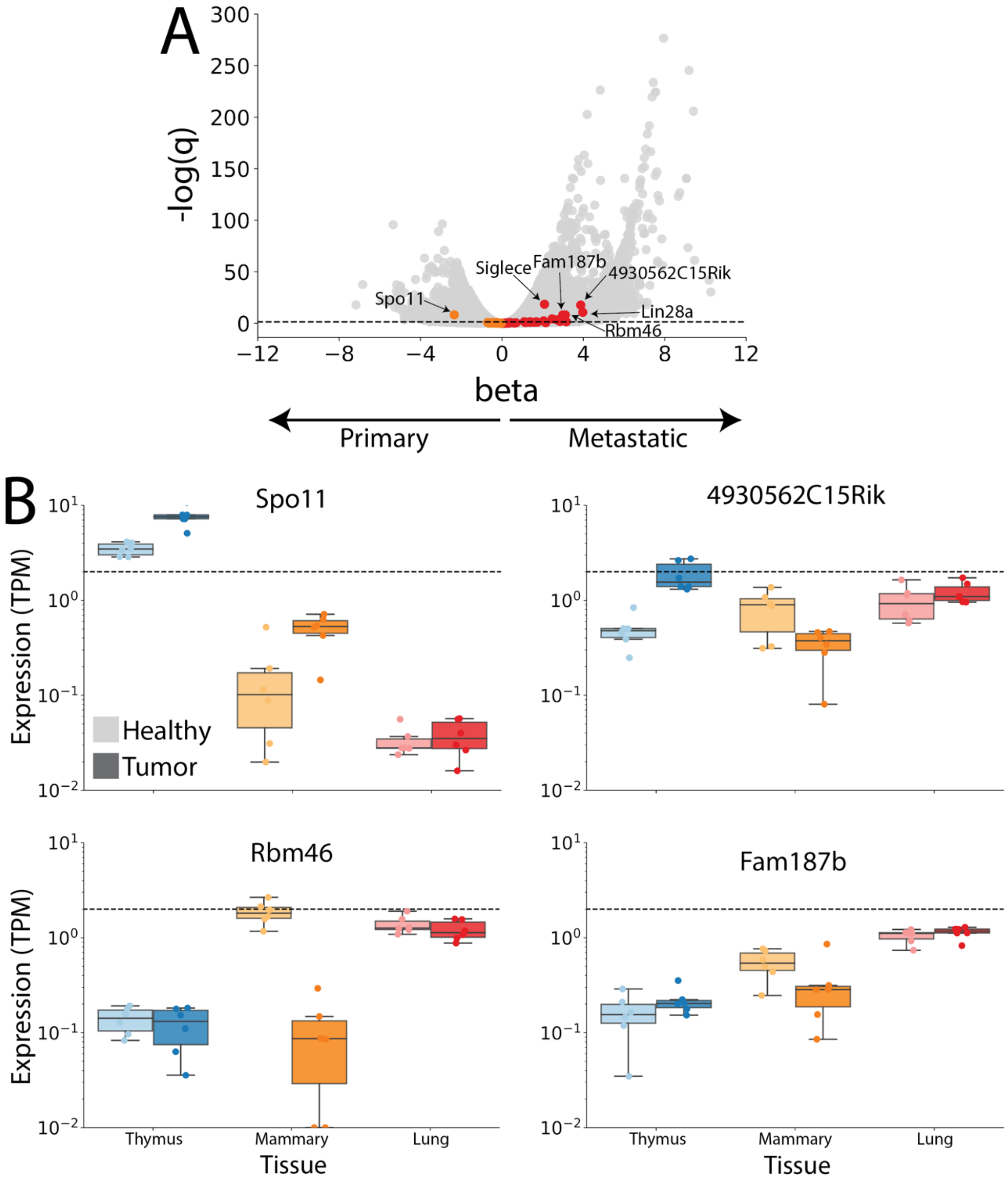


**Supplemental Figure 4. Differential expression between 4T1 primary mammary tumors and metastatic lung tumors in mice. (A)** Volcano plot showing differential expression between paired mouse primary mammary tumors and metastatic lung tumors. Homologs of human testis-exclusive genes are shown in color, with orange genes corresponding to enrichment in the primary tumor (beta<0) and red genes corresponding to enrichment in metastatic tumors (beta>0). **(B)** In addition to *Siglece* and *Lin28a* (shown in Figure 3E-F), expression patterns of the top differentially expressed genes (Spo11, 4930562C15Rik, Rbm46, Fam187b as indicated in panel (A)) are shown for thymus, mammary, and lung tissues taken from healthy (lighter color) and tumor-bearing (darker color) mice.

A

| | MHCI Score | MHCII Score | Total Score |
|---|---|---|---|
| L1 | 6.75 | 6.0 | 12.75 |

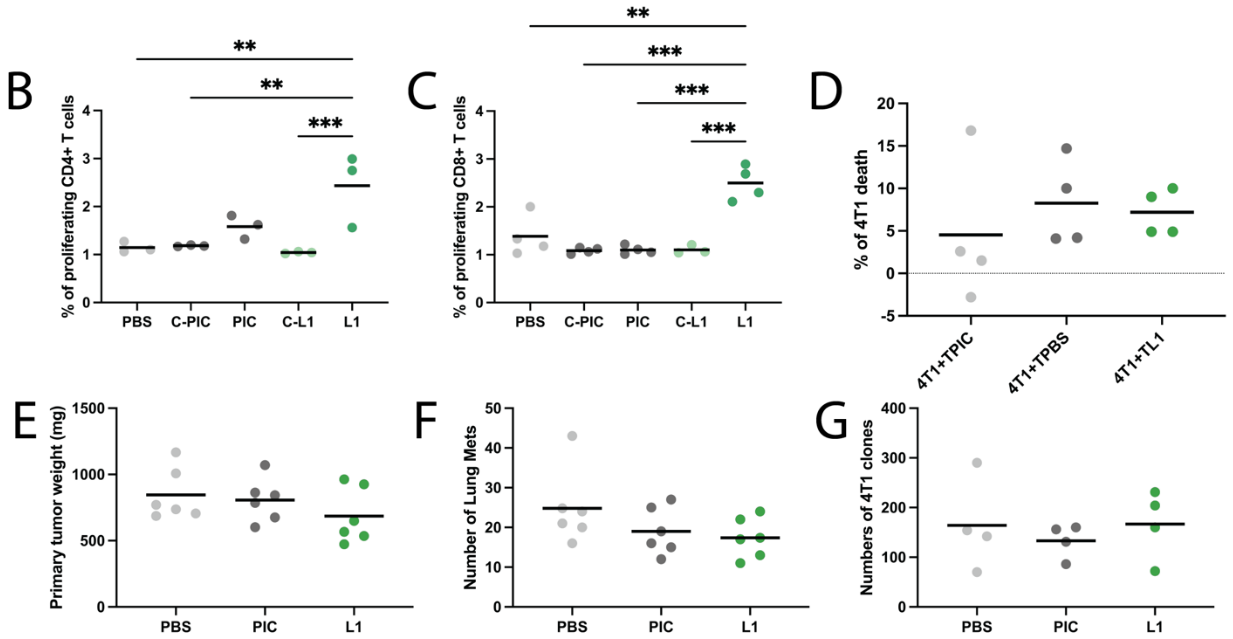


**Supplemental Figure 5. Efficacy of vaccination with Lin28a (L) based vaccine L1. (A)** Table showing predicted MHC binding scores, as outlined in Figure 3G, for the L1 peptide vaccine. **(B)** The L1 vaccine induced both CD4$^+$ and **(C)** CD8$^+$ T cell proliferation in pre-sensitized but not naïve control (c) mice as compared with poly(I:C) (PIC) and phosphate buffer solution (PBS) treated control mice. **(D)** As with L2 and expected given the lack of *Lin28a* expression in primary 4T1 mammary tumors, no difference was observed in 4T1 cell death following co-culture with tumor-cell infiltrates for mice vaccinated with L1 according to the preventative dosing strategy relative to PBS and PIC treated control mice. **(E)** Similarly, preventative dosing of the L1 vaccine did not decrease primary tumor size on day 28. **(F)** In contrast to L2, despite Lin28a expression in lung metastases, preventive treatment with the L1 vaccine did not reduce the number of lung metastases or **(G)** the number of lung 4T1 colonies. **$p<0.01$, ***$p<0.001$ by one-way ANOVA through Tukey's post hoc tests, all other pairwise comparisons not statistically significant.

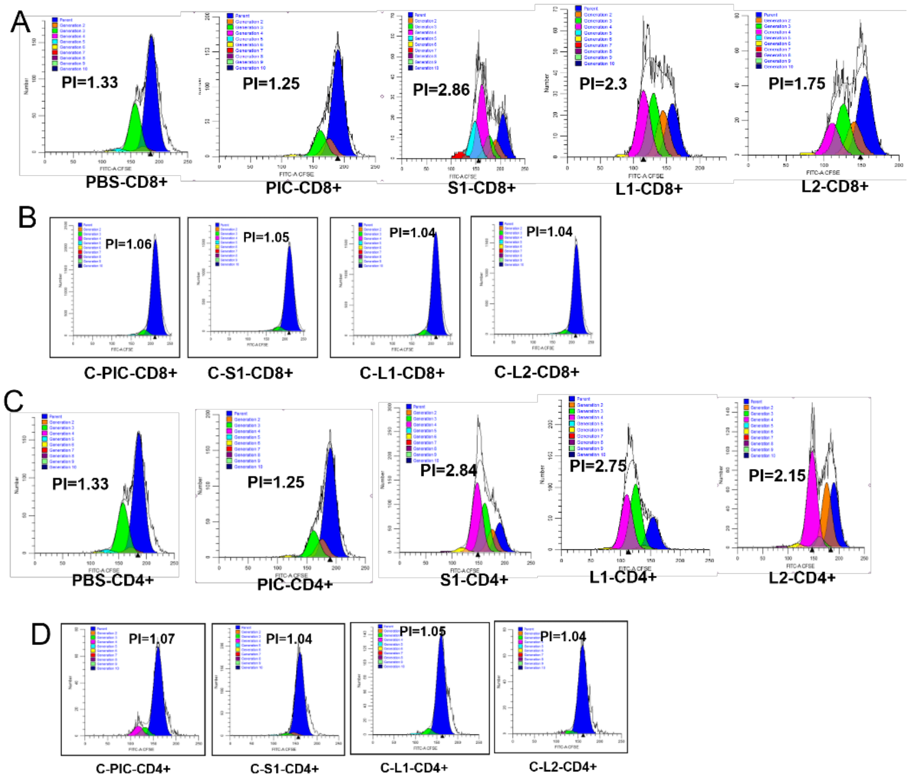


**Supplemental Figure 6. S1, L1, and L2 vaccine induced T cell proliferative responses. (A)** Flow cytometry data demonstrating proliferative response of sensitized CD8 + T cells towards S1, L1, or L2 peptides in pre-sensitized mice and **(B)** naive mice. **(C)** Proliferative response of CD4+ T cells in pre-sensitized and **(D)** naïve mice.

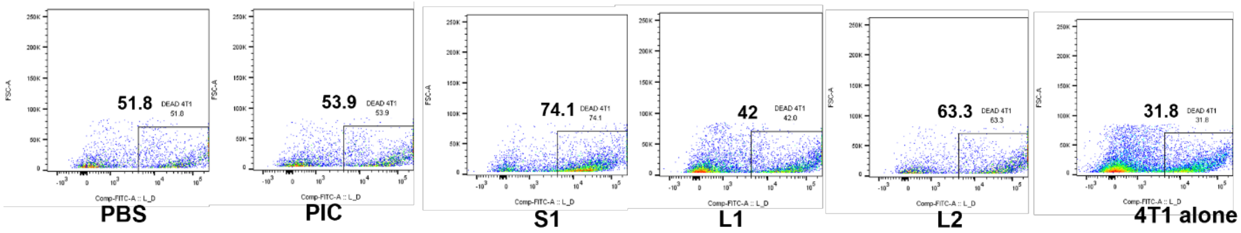


**Supplemental Figure 7. S1, but not L1 or L2, induces 4T1 cell death.** Representative flow cytometry plots showing the percentage of labeled 4T1 cell death following 24 hours of co-culture with vaccine or control treated tumor infiltrates.

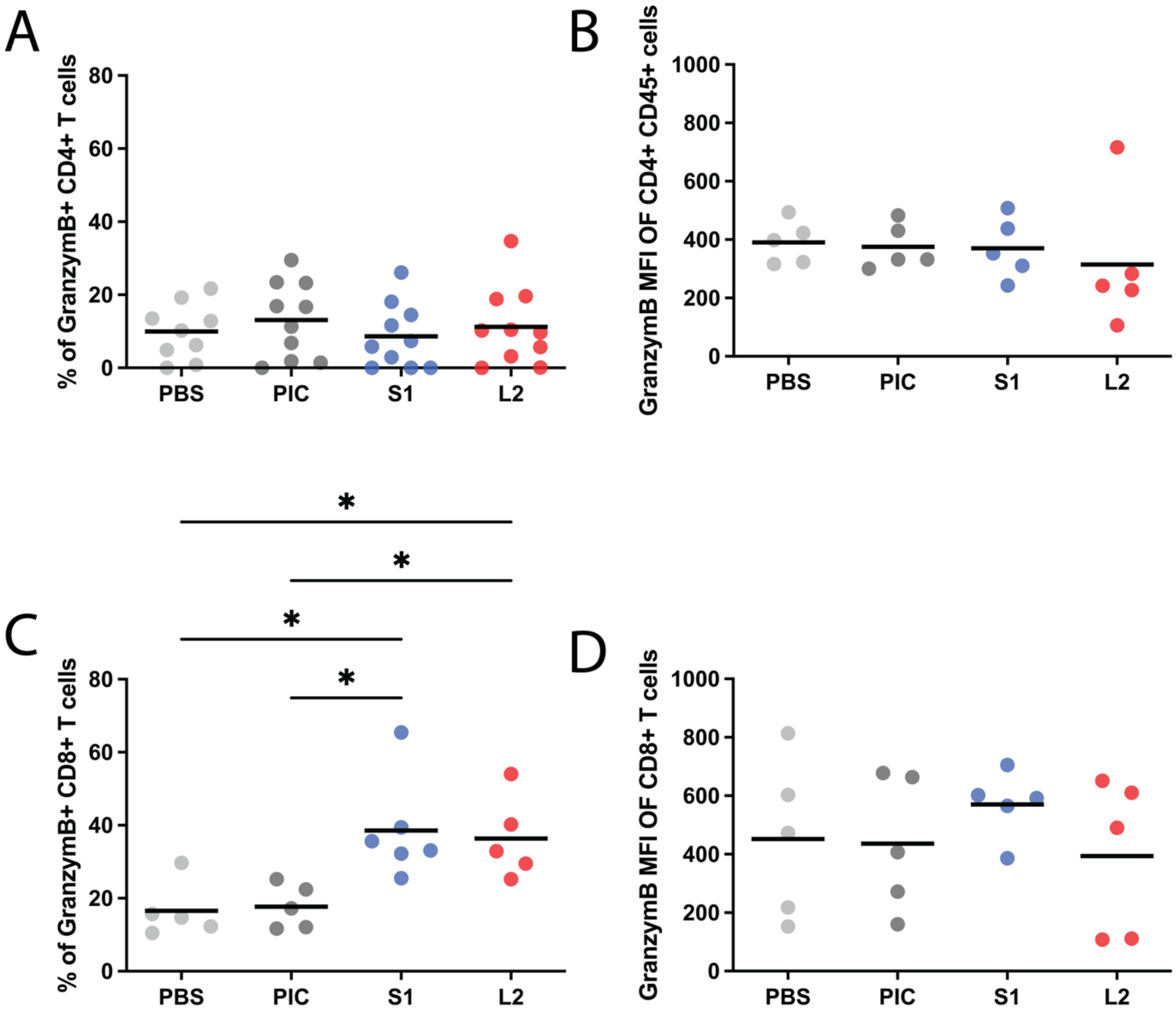


**Supplemental Figure 8. CT antigen vaccination stimulates CD8$^+$ expression of granzyme B. (A)** Preventative vaccination with S1 and L2 did not increase the fraction of CD4$^+$ T cells expressing granzyme B and **(B)** did not alter granzyme B expression level as represented by the mean fluorescent intensity (MFI). **(C)** Vaccination with both S1 and L2 increased the fraction of CD8$^+$ T cells expressing granzyme B, **(D)** but did not increase granzyme B expression in CD8$^+$ T cells. *$p<0.05$ by one-way ANOVA through Tukey's post hoc tests, all other pairwise comparisons not statistically significant.

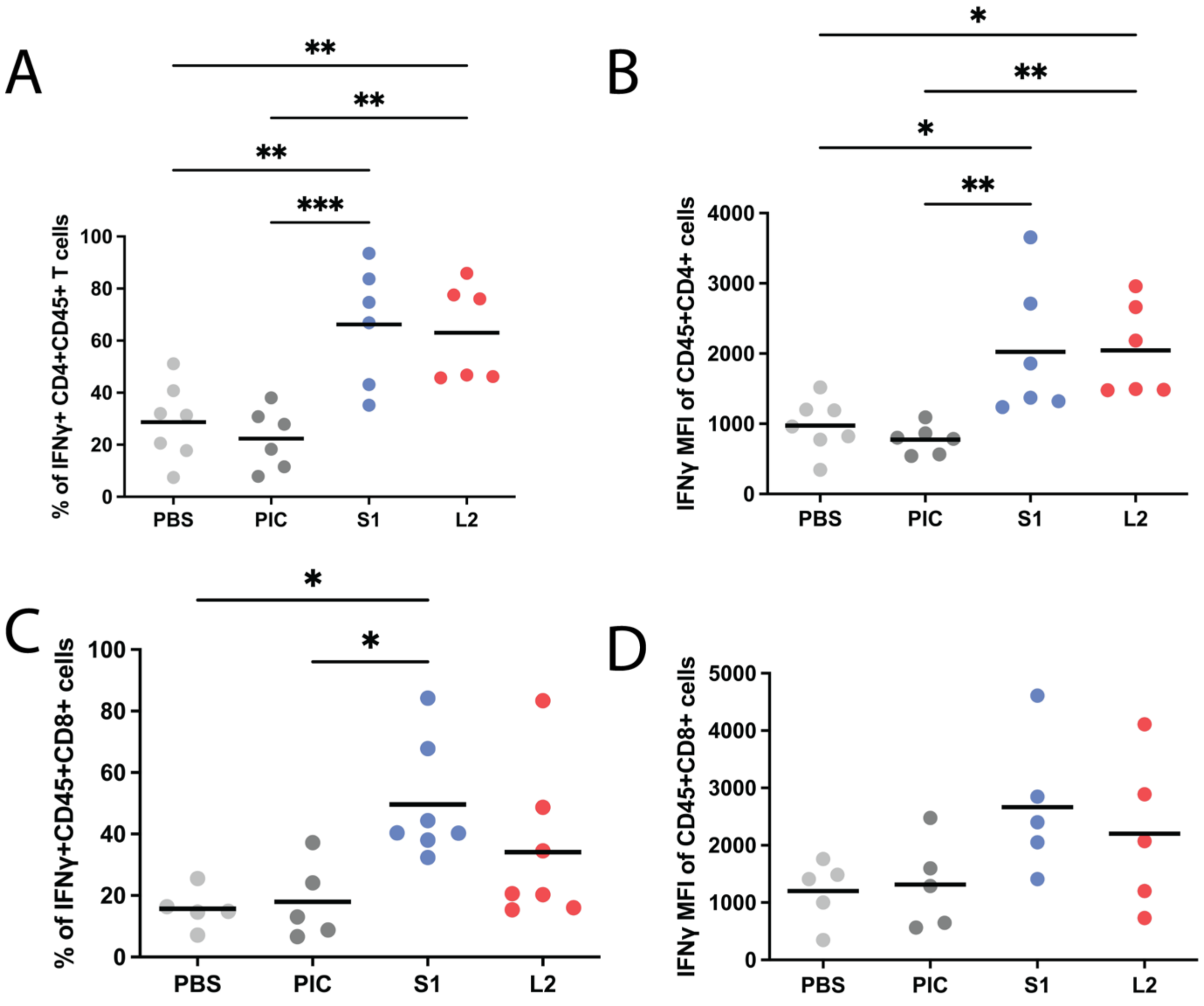


**Supplemental Figure 9 CT antigen vaccination stimulates CD4$^+$ and CD8$^+$ expression of interferon gamma (IFN$\gamma$). (A)** Preventative vaccination with S1 and L2 both increased the fraction of CD4$^+$ T cells expressing IFN$\gamma$ and **(B)** increased the average expression of IFN$\gamma$ within these CD4$^+$ T cells. **(C)** S1, but not L2, vaccination significantly increased IFN$\gamma$ in CD8$^+$ T cells **(D)** without increasing the average IFN$\gamma$ CD8$^+$ T cell expression. *p<0.05, **p<0.01, ***p<0.001 by one-way ANOVA through Tukey's post hoc tests, all other pairwise comparisons not statistically significant.
